# Disrupted Endothelial Cell Heterogeneity and Network Organization Impairs Vascular Function in Prediabetic Obesity

**DOI:** 10.1101/2020.05.07.083543

**Authors:** Calum Wilson, Xun Zhang, Matthew D. Lee, Margaret MacDonald, Helen H. Heathcote, Nasser M.N. Alorfi, Charlotte Buckley, Sharon Dolan, John G. McCarron

**Author notes:** **Subject terms**: Endothelium, Nitric Oxide, Vascular Biology, Vascular Disease. Author correspondence: John G McCarron or Calum Wilson, Strathclyde Institute of Pharmacy and Biomedical, University of Strathclyde, 161 Cathedral Street, Glasgow, G4 0RE, UK.

## Abstract

**Rationale:** Obesity is a major risk factor for diabetes and cardiovascular diseases such as hypertension, heart failure, and stroke. Impaired endothelial function occurs in the earliest stages of obesity and underlies vascular alterations giving rise to cardiovascular disease. However, the mechanisms that link weight gain to endothelial dysfunction are ill-defined. Increasing evidence suggests that, rather than being a population of uniformly responding cells, neighboring endothelial cells are highly heterogeneous and are organized as a communicating multicellular network that controls vascular function.

**Objective:** To investigate the hypothesis that disrupted endothelial heterogeneity and network-level organization contributes to impaired vascular reactivity in obesity.

**Methods and Results:** To study obesity-related vascular function without the complications associated with diabetes, we induced a state of prediabetic obesity in rats. Small artery diameter recordings confirmed nitric-oxide mediated vasodilator responses were dependent on increases in endothelial calcium levels and were impaired in obese animals. Single-photon imaging revealed a linear relationship between blood vessel relaxation and network-level calcium responses. Obesity did not alter the slope of this relationship, but impaired network-level endothelial calcium responses. The network itself was comprised of structural and functional components. The structural component, a hexagonal lattice network of endothelial cells, was unchanged in obesity. The functional network contained sub-populations of clustered agonist-sensing cells from which signals were communicate through the network. In obesity there were fewer but larger clusters of agonist-sensing cells and communication path lengths between clusters was increased. Communication between neighboring cells was unaltered in obesity. Altered network organization resulted in impaired, population-level calcium signaling and deficient endothelial control of vascular tone.

Specialized subpopulations of endothelial cells had increased agonist sensitivity. These agonist-responsive cells were spatially clustered in a non-random manner and drove network level calcium responses. Communication between adjacent cells was unaltered in obesity, but there was a decrease in the size of the agonist-sensitive cell population and an increase in the clustering of agonist-responsive cells

**Conclusions:** The distribution of cells in the endothelial network is critical in determining overall vascular function. Altered cell heterogeneity and arrangement in obesity decrease endothelial function and provide a novel framework for understanding compromised endothelial function in cardiovascular disease.

## Introduction

Obesity is a major risk factor for cardiovascular diseases such as hypertension and stroke^1-3^. Each of these diseases is precipitated by alterations in the endothelial cell lining of blood vessels^4^. The most frequently reported effect of obesity on endothelial function is a reduction in the ability of the cell layer to control vascular tone. Clinically, this endothelial dysfunction is indicated by reductions in hyperemia-induced forearm blood flow^5, 6^. Hyperemic flow is also impaired in obese children^7^, even in the absence of insulin resistance^8^, and short-term studies demonstrate reversible impairment of endothelium-dependent vasodilation with weight gain and loss^9, 10^. Thus, irrespective of age and in the absence of co-existing diseases, obesity is associated with impaired endothelial function.

Impaired endothelium-dependent function is also observed in various animal models of obesity, including obese Zucker^11, 12^ and JCR:LA-cp rats^13^, and rats fed a high-fat diet^14-16^. However, the mechanisms underlying endothelial dysfunction in obesity are unclear. Increased production of reactive oxygen species, which inactivates endothelial-derived nitric oxide (NO)^16, 17^, may explain reduced agonist-induced vasodilation in obese animals. Impaired endothelium-dependent vasodilation in obesity may also arise from changes in potassium channel activity^18-20^, NO release^20^, or how perivascular adipose tissue modulates endothelial function^21-23^. The number of diverse signaling mechanisms proposed to account for endothelial dysfunction in obesity demonstrates how complex changes occur in this cell layer, and point to changes in another regulator that is common to each dysfunction.

Intracellular Ca^2+^ is a second messenger that relays signals from numerous endothelial cell membrane receptors to effector proteins that govern vascular function^24^. For example, endothelial Ca^2+^ levels control endothelial derived hyperpolarizing factor, and regulate the synthesis and release of vasoactive mediators such as NO (reviewed ^25^). Ca^2+^ signals initiate when one or a few Ca^2+^ channels open to permit an influx of the ion into the cytoplasm. The resulting elevation in cytoplasmic Ca^2+^ can remain localized around the channel(s) or can grow into more global propagating Ca^2+^ waves by recruitment of neighboring channels. Studies of human arteries highlight a deficit in both local and global endothelial Ca^2+^ signals in obesity^26, 27^, but mechanistic insight into the disruption of local or global signaling is conflicting (local)^27, 28^ or absent (global).

Our recent work shows that endothelial control of vascular function is driven by cellular heterogeneity and large-scale, multicellular network Ca^2+^ dynamics^29^. Even within small vascular regions, endothelial cells are diversified and arranged in a network of signal detection sites^30^. Distinct clusters of cells are tuned to detect specific activators, and different clusters detect different stimuli. This arrangement allows the endothelium to detect and process multiple stimuli in parallel and to generate stimulus-specific responses^30-33^. These findings, together with studies that describe molecular^34-41^ and functional heterogeneity^40, 42^ in endothelial cells across vascular beds, and within single vessel segments, raise the possibility that dysfunctional vascular responses could arise from altered endothelial cell heterogeneity and disrupted network dynamics. Indeed, altered mulitcellular network behavior underlies disease development in a variety of physiological systems. For example, network dysfunction precedes the appearance of the earliest markers of neurodegenerative conditions^43, 44^, and contributes to islet failure in diabetes^45-47^. The role for endothelial heterogeneity in the pathophysiology of vascular disease is currently unknown.

Given the importance of endothelial function to the health problems in obesity and the absence of information on the pathophysiological correlate of endothelial cell heterogeneity, we investigated networked Ca^2+^ dynamics in large populations of endothelial cells in intact blood vessels. We show that agonist-evoked, endothelium-dependent vasodilation arises from network-level interactions that occur among a heterogeneous endothelial cell population, and that this network-level control is impaired in prediabetic obesity. Obesity alters the network, leading to a reduction in the size of the agonist-sensitive cell population, an increase in the clustering of sensory cells, and an increase the communication distance between cell clusters. These changes in endothelial cell network dynamics impairs the collective endothelial response and provides a new framework for understanding vascular dysfunction in obesity.

## Materials and Methods

All data underpinning this study is available from the authors upon reasonable request. An expanded Material and Methods section can be found in the Supplemental Materials.

### Animal models

All animal care and experimental procedures were conducted in accordance with relevant guidelines and regulations, with ethical approval of the University of Strathclyde Local Ethical Review Panel and were fully licensed by the UK Home Office regulations (Animals (Scientific Procedures) Act 1986, UK) under Personal and Project License authority. Twenty-eight male Wistar rats (9 weeks of age, weighing 349 ± 11 g) were split into two weight-matched groups and fed either a standard diet (SD group) or a high-fat diet (HFD group) for 24 weeks.

### Experimental techniques

First-order mesenteric arteries from SD- and HFD-fed rats were studied using pressure myography, and high-resolution single photon Ca^2+^ imaging of *en face* artery preparations with and without simultaneous assessment of vascular tone. For pressure myograph experiments, we used the open source pressure myograph system, VasoTracker^48^ and recorded outer vessel diameter in response to various treatments at 70 mmHg and 37°C. For Ca^2+^ imaging experiments, arteries were opened longitudinally, and endothelial cells were preferentially loaded with the Ca^2+^ indicator, Cal-520/AM (5 μM). Network-level Ca^2+^ signaling was imaged using a high (0.8) numerical aperture 16X objective (∼0.8 µm^2^ field of view). This 16X objective also permitted quantification of vascular contractility in opened arteries using edge-detection algorithms^49^. Subcellular Ca^2+^ signaling was imaged using a high (1.4) numerical aperture 100X objective. Network-level and subcellular Ca^2+^ signaling was assessed using custom Python software^30, 50^. Ca^2+^ responses were evoked by agonists or photolysis of caged inositol triphosphate (IP_3_).

### Statistics and data analysis

Summary data are presented in text as mean ± standard error of the mean (SEM), and graphically as mean ± SEM or individual data points with the mean indicated. Paired data points in plots are indicated by connecting lines. Concentration-response data were computed according to a three-parameter dose– response model and compared using two-way ANOVA with Tukey’s post-hoc test. All other data were analyzed using paired t tests, independent 2-sample t tests (with Welch’s correction as appropriate), ordinary two-way ANOVA with multiple comparisons, or repeated measures two-way ANOVA with multiple comparisons as indicated in the respective figure or table legend. All statistical tests were two-sided. A p value of < 0.05 was considered statistically significant.

## Results

### Impaired endothelium-dependent vasodilation in obesity

Rats on a high-fat diet developed obesity, without complications associated with diabetes (Figure S1). To examine endothelial function in this model of obesity, we first assessed endothelium-dependent vasodilation to acetylcholine (ACh). ACh evoked concentration-dependent relaxations of established tone (PE, 300 nM – 1 µM) in mesenteric arteries from SD and HFD rats (Figure 1A). In each group (SD & HFD), ACh-induced relaxations were inhibited by the selective M_3_ muscarinic receptor antagonist, 1,1-dimethyl-4-diphenylacetoxypiperidinium iodide (4-DAMP, 1 μM), or the NO synthase inhibitor, L-N^G^-Nitro arginine methyl ester (L-NAME, 100 μM), each applied intraluminally (Figure 1B-C, Figure S2 and Table 2). The inhibition of ACh-evoked relaxation by L-NAME was slightly enhanced by the additional presence of the K^+^ channel blockers TRAM34 (1 μM) and apamin (100 nM; Table 2, Figure S2) in SD and HFD groups. These results suggest that the major mechanism underlying relaxation is endothelial NO production.

**Figure 1.**
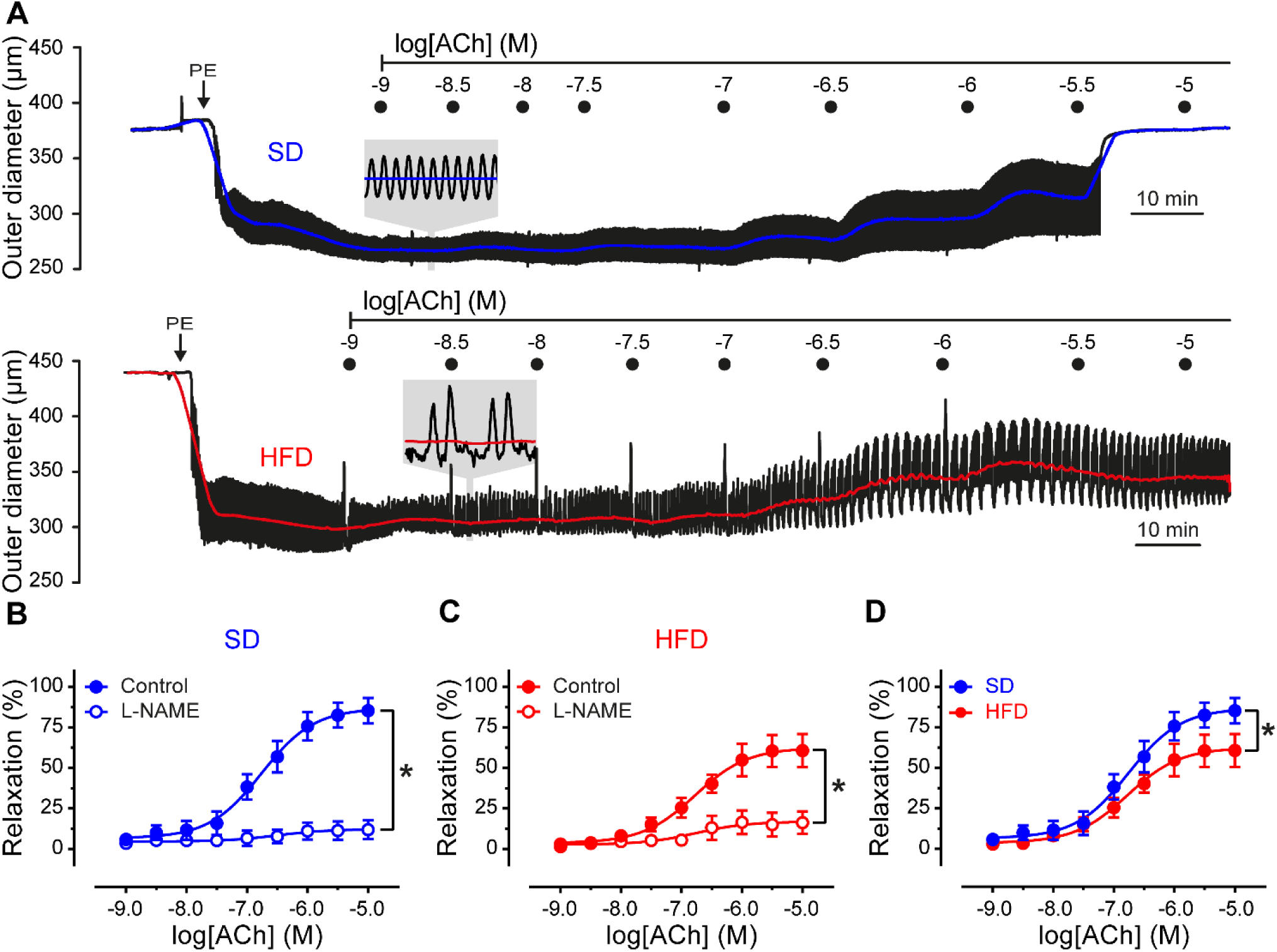
A high-fat diet impairs nitric oxide-mediated vasodilation. A) Representative mesenteric artery outer diameter traces in response to increasing concentrations of ACh. PE was continuously superfused and ACh was delivered intraluminally using a 10 cm H_2_O pressure gradient (∼ 200 µl min^−1^). B-D) Summary of diameter data comparing the concentration-dependent responses of arteries, from rats fed a standard diet (SD, blue) and a high-fat diet (HFD, red), to intraluminal ACh in the absence (filled circles) and presence (open circles) of the nitric oxide synthase inhibitor, L-NAME (100 µM). Data are mean ± SEM (n = 7 to 9). *p < 0.05, by comparison of sigmoidal fit parameters (top of sigmoid) using two-way ANOVA with Tukey’s post hoc test. Additional data shown in Figure S1 and tabulated in Table S2.

The muscarinic-receptor mediated, NO-dependent relaxations were impaired in mesenteric arteries from the HFD group (Figure 1D). However, there was no change in relaxation to the endothelium-independent NO donor, sodium nitroprusside (SNP, 10 μM; Table 2), suggesting that the processes involved in the production of NO are impaired in HFD-fed rats.

Endothelial production of NO is generally considered to be a Ca^2+^-dependent process ^25^. To determine if ACh-induced, NO-mediated vasodilation required an increase in endothelial Ca^2+^, we measured endothelial Ca^2+^ levels and vasoactivity before and after block of NO production or buffering endothelial Ca^2+^. Inhibiting NO synthesis using L-NAME (100 µM) enhanced PE-evoked contractions and inhibited ACh-evoked relaxations but had no effect on endothelial Ca^2+^ levels in SD or HFD groups (Figure S3 and Table 3). Preventing endothelial Ca^2+^ changes, by buffering Ca^2+^ with BAPTA, reduced basal intracellular Ca^2+^ levels, potentiated PE-induced contractions and inhibited ACh-induced relaxations in SD- and HFD-fed groups (Figure S3 and Table 4). The latter results demonstrate that endothelial Ca^2+^ signaling is required for endothelium-dependent relaxations.

### Population-level dysfunction of endothelial Ca^2+^ signaling in obesity

To examine the precise relationship between endothelial Ca^2+^ signaling and endothelium-dependent relaxation, we examined population-wide Ca^2+^ activity using high spatiotemporal resolution, wide-field single photon imaging (50-100 cells). We used an ACh concentration series to activate the cells while Ca^2+^ activity was recorded. The Ca^2+^ responses obtained in these experiments were normalized to maximal responses induced by ionomycin (1 µM), which was statistically similar in SD and HFD (Figure 2A, inset) and compared to the extent of relaxation at the same ACh concentration. As shown in Figure 2 there was a positive relationship between the amplitude of endothelial Ca^2+^ responses and relaxations evoked by ACh. In SD and HFD groups, the slope was linear and the gradient unity, suggesting a one-to-one relationship between global endothelial Ca^2+^ levels and vasodilator responses. Since the relationship between endothelial Ca^2+^ and relaxation was unchanged in obesity while relaxation was impaired, these findings suggest that endothelial dysfunction arises from an inability of the endothelium to generate a robust Ca^2+^ response in obese rats.

**Figure 2.**
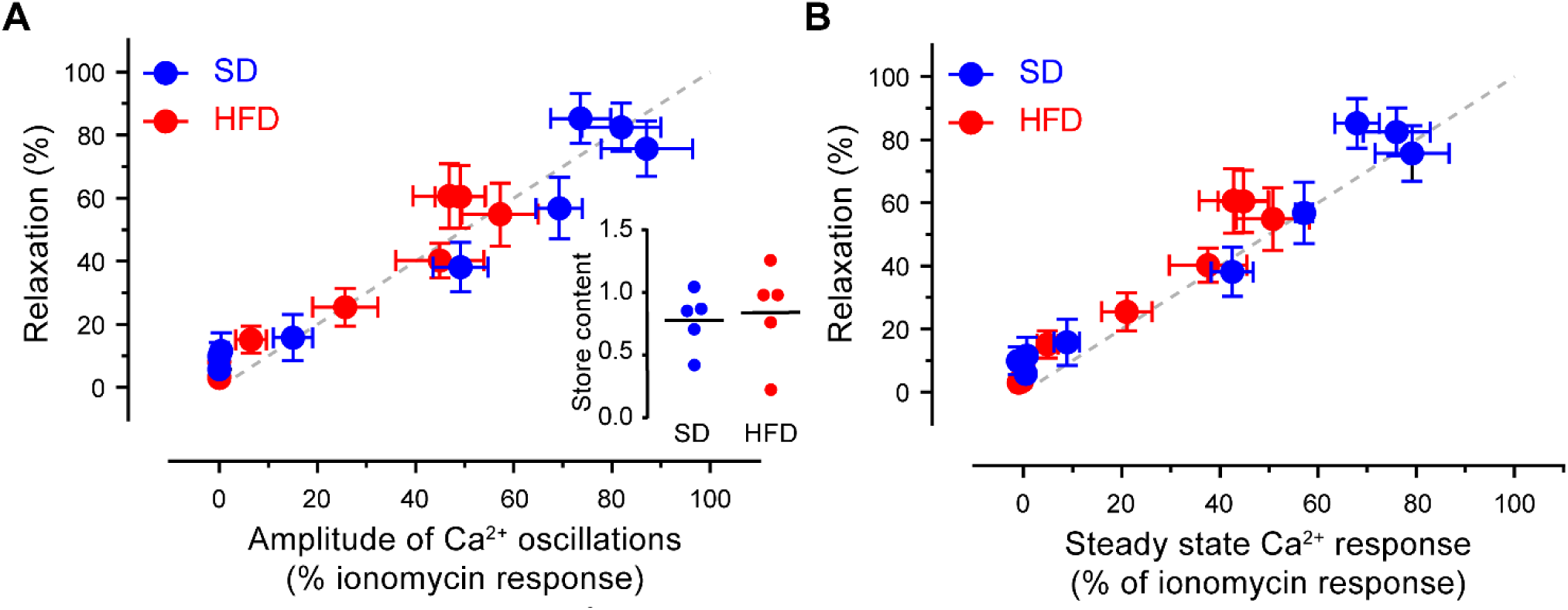
ACh-evoked endothelial Ca^2+^ levels and vascular relaxation are linearly proportional. A-B) Relationship between the amplitude of Ca^2+^ oscillations (A) or steady-state Ca^2+^ levels (B) and blood vessel relaxation from rats fed a standard diet (SD, blue) or a high-fat diet (HFD, red). The grey line represents a theoretical 1:1 correspondence between the Ca^2+^ parameter (x-axis) and relaxation. Inset in A plots the Ca^2+^ store content (area under the curve of ionomycin-evoked Ca^2+^ response) for SD and HFD groups.

To explore the mechanisms underlying the impaired Ca^2+^ response in obesity, we first determined the source of endothelial Ca^2+^ signals in our experimental model. To do this, the effects of several pharmacological interventions on various endothelial Ca^2+^ signaling parameters (number of activated cells, magnitude of the initial response, and magnitude of the response over 60 seconds) were examined (Figure 3 and Tables S5-S6). In each experimental group (SD or HFD), ACh-evoked (100 nM) Ca^2+^ responses were inhibited significantly by the selective muscarinic M_3_ receptor antagonist, 4-DAMP (1 µM), consistent with previous studies demonstrating the role of the M_3_ receptor in the endothelial response to ACh ^51^. In the absence of external Ca^2+^ (Ca^2+^-free PSS with1 mM EGTA included in PSS), endothelial cells remained responsive to ACh, although the rise in cytoplasmic Ca^2+^ concentration was not sustained (Figures S4). This result suggests that the initial response involved Ca^2+^ release from the internal store and that Ca^2+^ influx is required for the sustained phase. The phospholipase C inhibitor, U73122 (2 μM), but not its inactive analogue, U73343 (2 μM), prevented ACh-induced endothelial Ca^2+^ signaling. Similarly, two IP_3_ receptor (IP_3_R) inhibitors, 2-aminoethoxydiphenyl borate (2-APB, 100 μM) and caffeine (10 mM), each inhibited ACh-evoked increases in Ca^2+^. The broad-spectrum transient receptor potential channel antagonist, ruthenium red (RuR, 5 μM), did not reduce ACh-evoked endothelial Ca^2+^ activity. These results demonstrate that ACh evokes IP_3_-mediated Ca^2+^ release from the internal store, and suggest that the impaired response in obesity arises from either reduced Ca^2+^ release from activated IP_3_ receptors or a reduction in the ability of IP_3_ to generate Ca^2+^ release.

**Figure 3.**
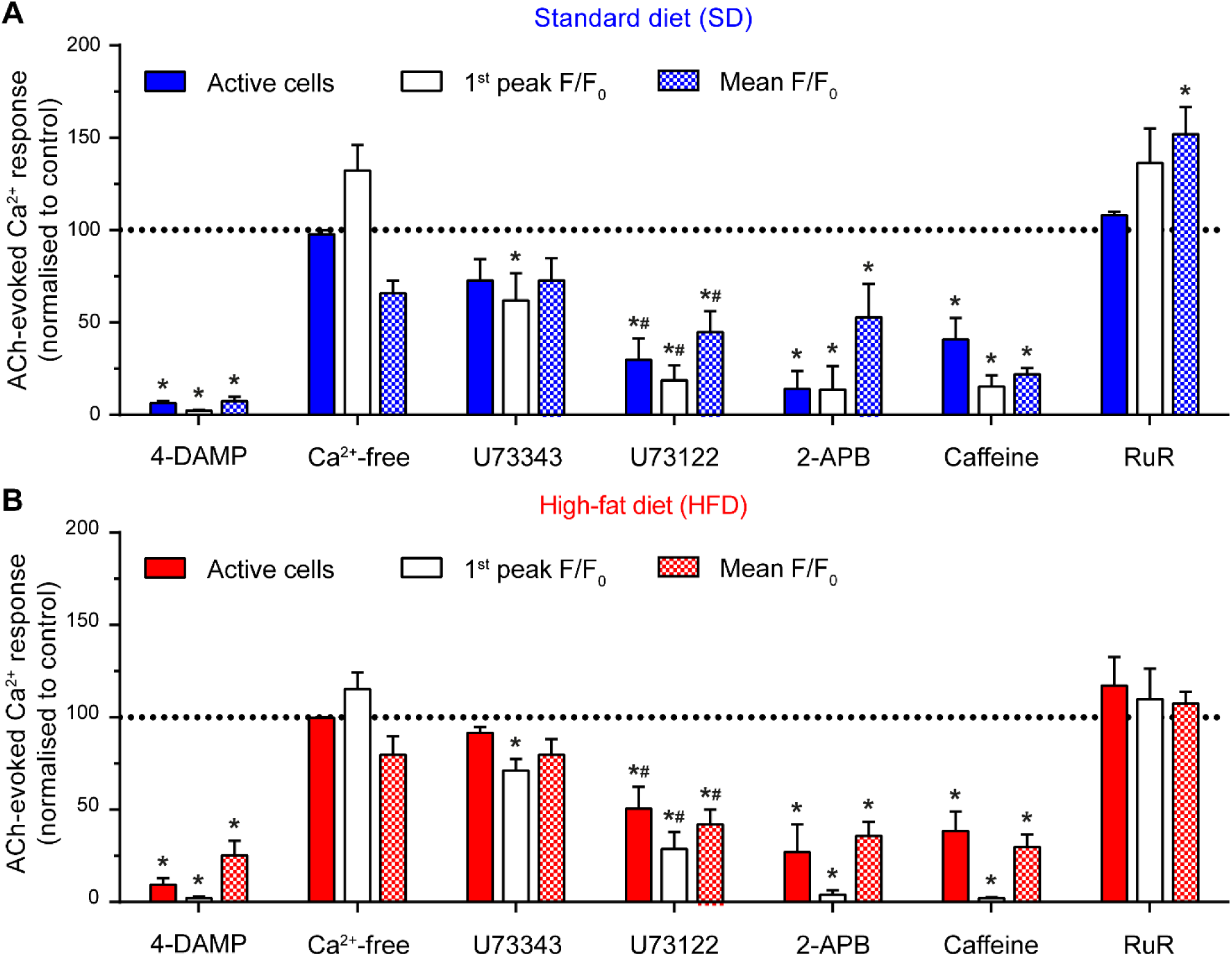
Acetylcholine activates endothelial Ca^2+^ signaling through the M_3_-PLC-IP_3_R pathway. Bar graph summarizing the effects of various pharmacological interventions on ACh (100 nM)-induced Ca^2+^ signaling in the endothelium of mesenteric arteries from rats fed either the standard diet (SD, A) or the high-fat diet (HFD, B). Ca^2+^-free PSS contained 1 mM EGTA. Concentrations of pharmacological inhibitors were: 4-DAMP (1 µM); U73343 (2 μM); U73122 (2 μM), 2-APB (100 μM); caffeine (10 mM); RuR (5 μM). All data are mean ± SEM expressed as a percentage of the control response in the same artery (response to ACh prior to pharmacological intervention). * indicates p < 0.05 versus corresponding control, # indicates p < 0.05 versus corresponding U73343, using repeated-measures two-way ANOVA with Tukey’s or Sidak’s multiple comparison test, as appropriate. Data normalized to control for presentation. All statistical analyses used raw data. Data tabulated in Tables S3-S4.

To investigate the mechanisms underlying alterations in IP_3_-mediated Ca^2+^ release in obesity, dynamic concentration-dependent activity from individual endothelial cells in intact blood vessels was examined. Spatial Ca^2+^ activity maps, generated from ΔF/F_0_ datasets show heterogeneity in the response of individual endothelial cells to ACh (Figure 4). ACh-responsive cells were not uniformly distributed but scattered across the endothelium in clusters. Increasing ACh concentrations activated a progressively larger percentage of the endothelial cell population (Figures 4A and S5A, top row). The concentration dependence of endothelial cell recruitment (% cells responding) was similar in the endothelium of SD and HFD groups (Figure 4B and Table S7). These results show heterogeneity in the endothelial response in SD and HFD rats.

**Figure 4.**
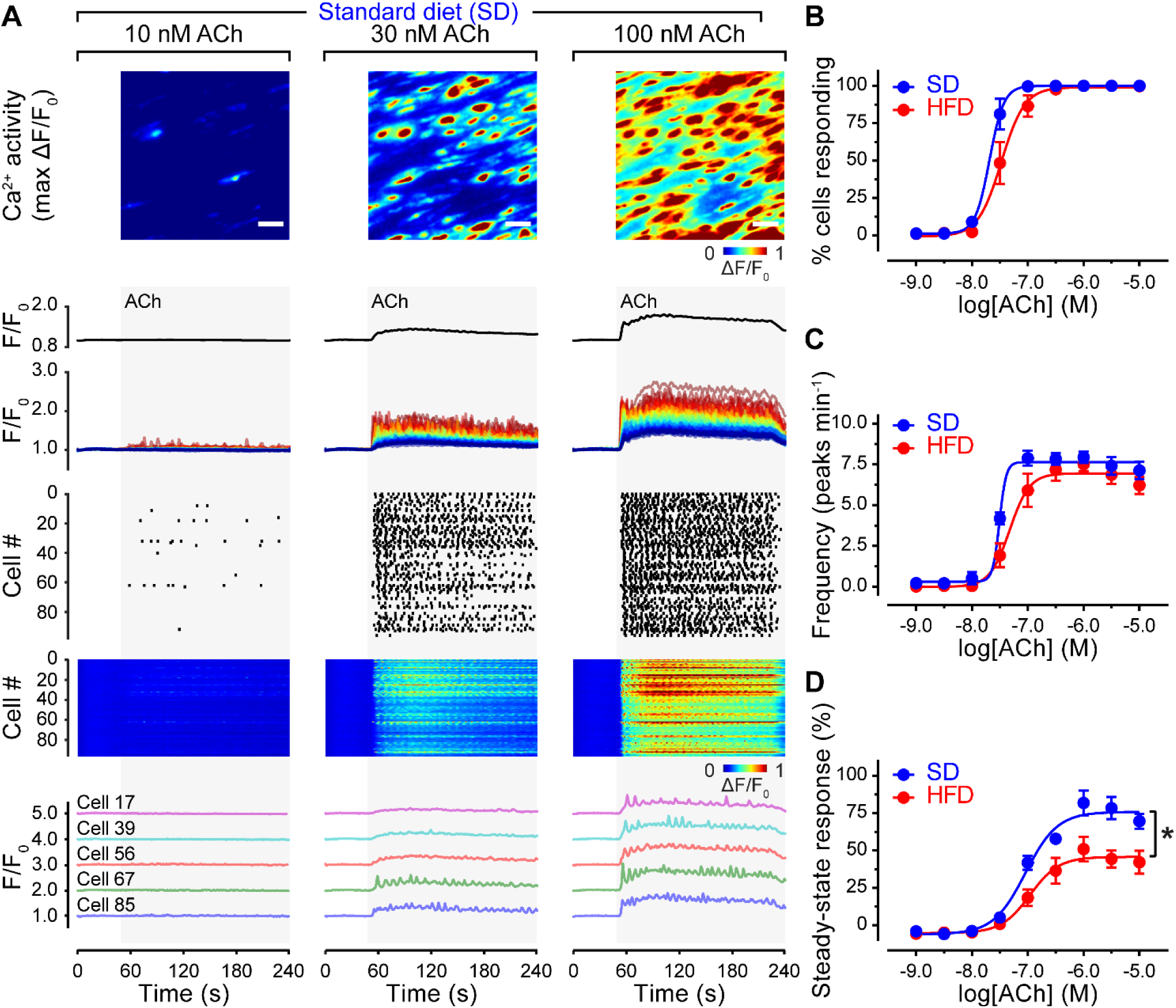
A high-fat diet impairs endothelial Ca^2+^ signaling. A) Concentration-dependence of ACh-evoked endothelial cell Ca^2+^ activity. The top row displays pseudocolored ΔF/F_0_ maximum intensity projections (shows all activated cells) of a single field of mesenteric endothelial cells (SD group) stimulated with the indicated concentrations of ACh. Scale bars = 20 µm. The second row displays the average Ca^2+^ signal across the field of view, whilst the third shows Ca^2+^ signals extracted from each cell in the field-of-view. Ca^2+^ traces are color coded according to the amplitude of the response to 100 nM ACh. The fourth and fifth rows are rastergrams (fourth) and heatmaps (fifth), each indicating spiking Ca^2+^ activity. The bottom row shows example traces from five separate cells. B-D) Summary of Ca^2+^ imaging data illustrating the concentration-dependence of the percentage of cells activated by ACh (B), the oscillation frequency (C), and the average level of the Ca^2+^ response of SD (blue) and HFD (red) rat endothelium. Data are mean ± SEM (n = 5 per group). * indicates significance (p < 0.05) by comparison of sigmoidal fit parameters (top of sigmoid) using two-way ANOVA with Sidak’s multiple comparison test. HFD data shown in Figure S3 and SD and HFD data tabulated in Tables S5-S6.

Temporal profiles of Ca^2+^ signals in individual cells (Figures 4A and S5A, bottom rows) also show the response to ACh is heterogeneous across responding cells, and dependent on agonist concentration. To quantify ACh-evoked Ca^2+^ activity, we measured the steady-state Ca^2+^ level, the magnitude of Ca^2+^ oscillations, and the frequency of Ca^2+^ oscillations in each cell. The magnitude (steady-state response, amplitude of oscillations) of Ca^2+^ increases evoked by ACh were significantly impaired in the HFD when compared to the SD controls (Figures 4D and S5C-D, and Table S7). However, there was no significant difference between the concentration-response relationship for Ca^2+^ signal oscillation frequency in HFD when compared to SD-fed rats (Figure 4C and Table S7). Thus, in obesity although endothelial cells remain able to encode information in the frequency of Ca^2+^ signals, the amplitude of this activity is impaired.

One possible explanation for these results is that IP_3_ itself is less efficient in evoking Ca^2+^ release from internal stores in obesity. To examine this possibility, we bypassed PLC-dependent IP_3_ production using photolyzed caged-IP_3_ to directly activate IP_3_ receptors. Elevations in Ca^2+^ evoked by photolysis of caged-IP_3_ were similar in the SD and HFD groups (Figure S6). This finding rules out the possibility that the reduced ACh-evoked endothelial Ca^2+^ activity in the HFD group arises from an impaired ability of IP_3_ to evoke Ca^2+^ release.

### Abnormal endothelial Ca^2+^ responses reflect altered endothelial cell heterogeneity

The results presented so far suggest that obesity impairs vascular responses by reducing endothelial Ca^2+^ signaling, and that an inability of IP_3_ to evoke Ca^2+^ release does not explain the findings. Intracellular communication is critical in determining coordinated endothelial responses. Interactions among endothelial cells permit Ca^2+^ signals in discrete clusters of cells to coalesce into networked signaling patterns of activity that drive tissue-level responses ^52, 53^. Thus, we hypothesized that the endothelium may be unable to generate robust population-level Ca^2+^ responses in obesity because of either or both of: 1) alterations in the spatial distribution of endothelial sensitivity to ACh; and 2) altered communication between endothelial cells.

To test these possibilities, we examined the organization, functional relationships, and patterns of IP_3_-mediated Ca^2+^ activity occurring between neighboring endothelial cells (Figure 5). In these experiments, large areas of endothelium (∼0.8 mm^2^) were imaged and regions of interest were generated for each cell visualized (3667 cells, n = 6 for SD; 3051 cells, n = 6 for HFD). In both experimental groups, endothelial cells were arranged in a lattice network and each cell possessed an average of six immediate neighbors (Figure 5A-C). This physical structure was unchanged in obesity. However, the structural connections among cells might give rise to altered functional connections in obesity. As a first step in examining the functional network in control and obesity, we identified cells that were unambiguously sensitive to ACh – i.e. those cells that responded to ACh before any immediate neighbor (ACh-sensitive cells; Figure 5D&F). There was a smaller density of these ACh-sensitive cells in the HFD group compared to the SD group (Figure 5G), and this generated an increased distance (path length) between ACh-sensitive cells (Figure 5H). Thus, a significant change in endothelial responsiveness in obesity arises from a decreased density of agonist sensing cells.

**Figure 5.**
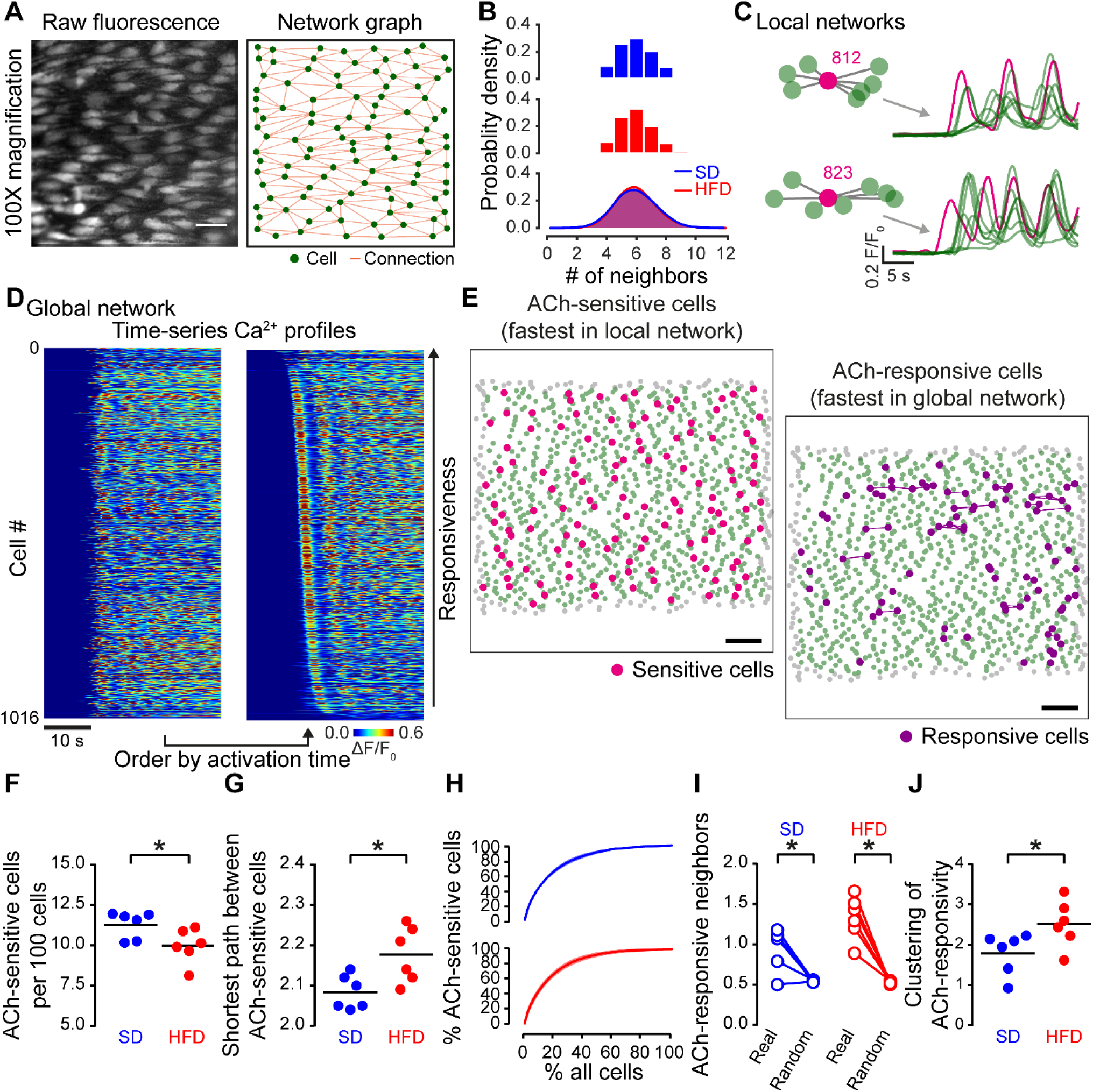
High-fat diet alters endothelial cell heterogeneity. A) High-resolution image of the endothelium (left) and corresponding structural network (right). Network connectivity was computed using line segments regions-of-interest. Green circles indicate the centroid of each endothelial cell, orange lines indicate connections between adjacent cells. B) Probability distribution of endothelial cell connectivity showing the number of neighbors each cell has. C-E) Endothelial heterogeneity was assessed using two methods. First, local cell networks were interrogated to reveal ACh-sensitive cells (those that respond to stimuli before any neighbor; C and E). Endothelial cells were also ranked (globally) by the time of their response to ACh to reveal the top 10% most ACh-responsive cells (D and E). F-G) Summary data showing the effect of a high-fat diet on the density of ACh-sensitive cells (F) and the mean distance (number of cells) between ACh-sensitive cells (G). F) Graph showing that ACh-sensitive cells tend to be the first cells to respond. I-J) Summary data showing the clustering of ACh-responsive neighbors compared to an equivalent random model (I), and a comparison of clustering between SD and HFD groups (J). * indicates statistical significance (p < 0.05) using paired t-test (I) or unpaired t-test (F, G, J) with Welch’s correction as appropriate. Scale bars = 20 µm (A) or 100 µm (E).

To determine if endothelial cell clustering was altered in obesity, we next examined the distribution of agonist-responsive cells. In this analysis, we ordered cells by the speed at which they responded to ACh and identified the first 10% of cells to respond as ACh-responsive cells (Figure 5E&F). Ca^2+^ responses in this fast-responding population of cells are likely due to direct agonist activation, rather than indirect activation arising from signals originating in neighboring cells and contained a large percentage of ACh-sensitive cells (Figure 5F). In control and obesity, ACh-responsive cells had more ACh-responsive neighbors than would be expected if the endothelial cells were randomly distributed with respect to responsivity (Figure 5J), i.e., ACh-responsive cells are clustered throughout the endothelial network^30, 33^. Furthermore, compared to control, clustering of the ACh-responsive cell population was increased in obesity (Figure 5K). This observation is significant as the increased clustering in HFD decreases the ability of responsive cells to engage with and recruit unresponsive cells across the network.

We next examined endothelial network communication by determining the functional connectivity of adjacent cells. In this analysis, a pairwise cross-correlation analysis was used to assess signal similarity between all neighboring endothelial cell pairs (i.e. quantifying the level of shared Ca^2+^ activity with time, Figure S7). This type of analysis provides information on how cells encode information at the population level and how they interact, yielding insights into network communication mechanisms. In SD and HFD-fed animals, the mean pairwise correlation coefficients significantly exceeded chance levels (Figure S7E). However, the extent of pairwise synchronicity was statistically similar in the two groups (Figure S7F). This result indicates that the extent of communication between activated cells is unaltered in obesity. In support of this conclusion, Ca^2+^ wave propagation initiated by focal release of IP_3_ was similar in SD and HFD (Figure S8). This finding also suggests that communication among cells is unaltered in obesity. Thus, whilst intercellular communication is unaltered, the density of ACh-sensitive cells is reduced, and the clustering of agonist-responding cells is increased in obesity.

### Local Ca^2+^ signaling

An alternative route by which the endothelial network may be compromised is via changes in local signaling events. In many vascular diseases (e.g. hypertension ^49^), dysfunctional endothelium-mediated responses arise from impaired signaling between endothelial and smooth muscle cells. Such signaling occurs via specialized myoendothelial projections (MEPs) that extend from endothelial cells, through holes in the internal elastic lamina (IEL), to smooth muscle cells. For example, IP_3_-evoked Ca^2+^ increases at MEPs may activate endothelial NO synthase and IK/SK channels ^54, 55^, which each promote vascular relaxation. Because of this, we tested if the impaired vasodilation identified in HFD-fed rats might also reflect disruptions in endothelial-smooth muscle cell signaling. First, we determined the degree of myoendothelial connectivity by imaging the IEL (Figure S9A-B). There was no clear change in IEL structure, as evidenced by similar IEL hole density, mean IEL hole size, and mean percentage area of IEL occupied by holes (IEL hole coverage; Figure S9C-E and Table S8). We next examined the relationship between spontaneous Ca^2+^ activity and IEL holes (Figure S10 and Table S8). In contrast to ACh-evoked Ca^2+^ activity, basal (unstimulated) Ca^2+^ events are mostly subcellular waves and the initiation site of individual events can be readily identified (see also ^29, 32^). Importantly, irrespective of diet, we found that endothelial Ca^2+^ events occurred closer to MEPs than would be expected than if they occurred randomly throughout the cytoplasm (Figure S10C-D and Table S8). However, there was no difference in the extent of endothelial Ca^2+^-event-MEP coupling in SD- and HFD-fed rats (Figure S10E and Table S8). Thus, it is unlikely that alterations in myoendothelial coupling contributes to the vascular dysfunction in obesity observed in the present study.

## Discussion

Obesity, a disease characterized by excess body fat, is associated with type 2 diabetes, metabolic syndrome and cardiovascular diseases such as hypertension and stroke. Impaired endothelial function is a hallmark of obesity-related cardiovascular diseases^4^, but mechanisms underlying the dysfunction are poorly understood. Here, we identify aberrant population-level activity in native endothelial cell networks of obese rats. Altered endothelial cell network circuits arose from compromised cell heterogeneity and increased clustering of sensory cells, and resulted in deficient encoding of vasoactive stimuli into population level Ca^2+^ responses. Functionally, this network deficit manifested as impaired vasodilator responses. Abnormal vasodilator function occurred despite a lack of effect on intercellular Ca^2+^ signal propagation or pairwise synchrony between adjacent endothelial cells, suggesting a spatial network disruption rather than failure in network communication itself. Together, these findings: 1) highlight the role of population level interactions in governing endothelial function, 2) offer functional insight into endothelial cell heterogeneity seen in many genetic studies; and 3) provide support for a new cellular heterogeneity and organization hypothesis of endothelial dysfunction in cardiovascular disease.

Endothelial cells are interlaced with one another to form a regular mesh (hexagonal lattice) network in which each cell has ∼6 adjacent neighbors. This structural architecture is unaltered by obesity. However, recent application of network theory to endothelial cell populations has started to uncover how rich functionality emerges from these structural connections and how network connectivity permits multiple physiological processes to be controlled^30-33^. Observing agonist-mediated endothelial Ca^2+^ activity at the population level reveals spatial modules, or clusters of sensory cells, that detect activators and recruit agonist-insensitive cells to drive network activity. Recruitment occurs via the transmission and propagation of Ca^2+^ signals and increases the overall population of active cells contributing to the vascular response. Here, we show that the size of the sensory cell population is reduced in obesity and there is an increase in the clustering of agonist-responsive cells, i.e., there are fewer but larger modules. Increased clustering of sensory cells results in activated cells communicating to a greater extent with other agonist-activated cells. Since endothelial Ca^2+^ signals decay with transmission distance^56-58^, the increased clustering limits recruitment of agonist-insensitive cells. Thus, changes in endothelial cell distribution disrupt collective endothelial cell behavior, reduce population-level Ca^2+^ responses and so impair endothelial vasodilator responses.

Endothelial cells in different vascular beds are heterogeneous and endowed with functions suited to the organs they vascularize^59^. Endothelial cell variation also occurs even within small segments of blood vessels. Intra-vessel endothelial heterogeneity has been measured in the distribution of a wide variety of proteins including α-adrenoceptors^60^, angiotensin II receptors^34^, cannabinoid receptors^60^, histamine receptors^61^, muscarinic receptors^30, 51, 61, 62^, purinergic receptors^30^ and von Willebrand factor^34^. The functional significance of endothelial cell heterogeneity is only beginning to emerge. As an example, a distinct subpopulation of aortic endothelial cells with self-renewal capacity drives tissue repair^63^. In some tissues, heterogeneous cell populations may organize into spatial domains with coherent gene expression^64^, and it is thought that such multicellular groups form microcircuits, the building blocks of information processing. Clustering of endothelial cells permits the distribution of activities across space, i.e., different functional units can perform different processes simultaneously^30-32^. Such parallel processing requires cells that are responding to one stimulus to be resilient to interference from neighboring cells and this is facilitated by the organization of cells into clusters. Clustering will also increase the concentrations of diffusible messengers (e.g. NO), so increasing their effective range, by overwhelming local breakdown mechanisms. Thus, network organization and cellular heterogeneity are fundamental to how the endothelium responds to external environmental drivers.

Here, we demonstrate that the relationship between population-level endothelial Ca^2+^ signaling and vasodilator function is linear and injective. How endothelial microcircuits interact and coordinate to control vascular reactivity via tissue-level endothelial Ca^2+^ signaling is an important question, particularly as the mapping of endothelial Ca^2+^ levels to vascular relaxation is unaltered in obesity. Indeed, the decrease in vasodilator function in obesity appears to be a direct consequence of a reduced ability of the endothelium to generate population wide increases in intracellular Ca^2+^, rather than any impairment of downstream signaling mechanisms. Cooperativity among cells is essential for generating coherent global responses and necessitates communication among endothelial cells. Such communication is achieved in endothelial cell networks via the transmission of IP_3_ or Ca^2+^ or both, through intracellular gap junctions that connect neighboring cells ^52, 65^. Two findings indicate that, in obesity, endothelial cells remain connected and retain the ability to communicate with each other. First, the extent of pairwise similarity between Ca^2+^ signals of neighboring cells was similar in control and obesity, suggesting comparable functional connectivity. Second, evoked Ca^2+^ wave propagation was similar in control and obesity. Consistent with a cell heterogeneity hypothesis of endothelial dysfunction, these findings suggest that the mechanisms underlying endothelial cell cooperativity are unaltered in obesity.

Whilst our study demonstrates that altered endothelial cell heterogeneity and network organization are responsible for deficient vasomotor control in obesity, we have not addressed which mechanisms are responsible for promoting altered endothelial cell clustering. Molecular and functional differences exist between endothelial cells of various blood vessels and vascular beds, and this diversity may arise due to microenvironmental factors. In support, transplanted endothelial cells gain structural phenotypes and gene expression patterns associated with new host tissue^66, 67^ and endothelial cells with distinct molecular signatures regress towards a common phenotype upon in vitro expansion^39, 68^. In obesity, differences in the local environment may arise from variation in lipid accumulation among endothelial cells^69^. Heterogeneity may also arise due to specific origins of endothelial cells rather than the local environment. Two distinct developmental origins give rise to endothelial cells in mature blood vessels^70^. Heritable factors from these different origins may explain why proliferative endothelial cells exist alongside cells with a lower proliferative potential^63, 71^, and give rise to clusters of endothelial cells with similar properties via clonal expansion^37, 38^. Interactions between heritable factors and local environmental conditions may also explain endothelium dysfunction in obesity. Clarifying exactly how functional endothelial cell heterogeneity emerges and is impacted in disease states such as obesity is a major area for future work.

In conclusion, the present results show that the endothelium is a collection of exquisitely organized cells, and that this organization governs endothelial function. Impairment of vascular function in obesity arises from alterations in endothelial cell heterogeneity, specifically the altered distribution of sensory cells, and results in deficient network-level function (vasodilation). As the mechanisms disrupting endothelial network function are not yet clear, significant work remains to address a heterogeneity model of vasomotor dysfunction. Future studies will investigate drivers of altered endothelial heterogeneity and network dysfunction in obesity and its related vascular diseases. As collective cell behavior also controls the regeneration of injured endothelium^71^, the consequences of network dysfunction are likely to extend far beyond the control of blood vessel diameter. Understanding the full functional significance of endothelial cell network organization will provide a deeper understanding of the pathophysiology of vascular disease.

## Acknowledgements

This work was funded by the Wellcome Trust (204682/Z/16/Z; 202924/Z/16/Z) and the British Heart Foundation (PG/16/54/32230; PG16/82/32439, PG/20/9/34859), whose support is gratefully acknowledged.

## Author Contributions

JGM & CW developed the concept. NMNA & SD developed the animal model. CW, XZ, MM, MDL, CB & HRH performed the experiments, and analyzed the data. CW, XZ, MDL, CB, HRH, & JGM interpreted the data. CW & JMG drafted the manuscript. All authors revised and edited the manuscript and approved the final version of the manuscript.

## Non-standard Abbreviations and Acronyms

2-APB: 2-aminoethoxydiphenyl borate
4-DAMP: 1,1-dimethyl-4-diphenylacetoxypiperidinium iodide
ACh: acetylcholine
ANOVA: analysis of variance
HFD: high-fat diet
IEL: internal elastic lamina
L-NAME: L-NG-Nitro arginine methyl ester
IP3: inositol triphosphate
MEP: myoendothelial projection
NO: nitric oxide
RuR: ruthenium red
SD: standard diet
SNP: sodium nitroprusside

