## Supplementary information for "Disrupted Endothelial Cell Heterogeneity and Network Organization Impairs Vascular Function in Prediabetic Obesity"

### Detailed Methods

#### Animals and nutritional management

Twenty-eight male Wistar rats (9 weeks of age, weighing 349 ± 11 g), purchased from Invigo (U.K.), were used in the study. Male rats are widely-used in experimental models of diet-induced obesity with a wealth of background information, and were used to aid interpretation of results. All rats were acclimatized to the animal housing facility for one week prior to the study and were housed in pairs or triplets (RC2F cages with Sizzle nesting material; North Kent Plastic, UK). Rats were maintained (temperature of 21°C ± 2°C, 45-65% humidity, 12-hour light-dark) with *ad libitum* access to a standard diet (SD; Rat and Mouse No.1 Maintenance, 801151, Special Diet Services, UK) and water.

At the end of the acclimatization period, fourteen rats were switched from the standard diet to a high-fat diet (HFD). Fourteen weight-matched rats continued to receive standard diet. The nutritional composition of the standard diet was: 66.7% calories from carbohydrate, 14.4% calories from protein and 2.7% calories from fat with a total calorific value of 14.74 MJ $kg^{-1}$ . The nutritional composition of the high-fat rodent diet (HFD; High Fat (M), 821424, Special Diet Services, UK) was: 44.9% calories from carbohydrate, 19.8% calories from protein, and 22% calories from fat with a total calorific value of 19.67 MJ $kg^{-1}$, respectively.

All animals were maintained on their respective diets for 24 weeks, at which point the study was ended. During this period, individual body-weight, and food/water consumption per cage were measured weekly. Blood glucose levels (non-fasting, tail-vein samples) were measured every second week using a hand-held glucometer (Alpha-Trak 2, Zoetis, USA). At week 20, an insulin tolerance test (ITT) was also performed on a subset of fasted rats (4-6 hours) from each experimental group. Blood glucose was measured from the tail vein, and insulin was then administered by intraperitoneal injection at a concentration of 1 U $kg^{-1}$. Blood glucose was measured 60 minutes after the insulin injection. At the end of the study, rats (non-fasted) were euthanized by intraperitoneal overdose of sodium pentobarbital (Euthatal, 200 mg $kg^{-1}$; Merial Animal Health Ltd, Woking, UK). Blood was obtained by cardiac puncture for measurement of plasma insulin, total triaglyceride (TAG), and total cholesterol using commercial kits (Rat Insulin ELISA kit (Thermo Scientific, UK); Total Serum Triglyceride Determination Kit, method B1 (Sigma-Aldrich, UK); Cholesterol Quantitation Kit (Sigma-Aldrich, UK). Of a total of twenty-eight animals, twenty-seven progressed to the end of the study. One rat from the control group died early.

**Body weight and metabolic parameters**

HFD-fed rats had significantly higher weight gain from week 4 onwards when compared to rats on a standard diet (SD) (Figure 1A-B; n = 14 for HFD, n = 13 for SD). By the end of the study, the mean increase in weight was 86 ± 4 % (from 349 ± 11 g to 654 ± 33 g) for HFD-fed rats and 58 ± 3 % (from 358 ± 16 g to 565 ± 26 g) for SD-fed rats (Table 1). Except for 5 weeks, energy intake was significantly higher in HFD-fed rats than in SD-fed rats throughout the study (n = 6 cages per group). Water intake was similar in the two groups (Figure 1E and Table 1).

At the end of the study, resting blood glucose concentrations in the HFD group were similar to those in the SD group (Figure 1F and Table 1). There was no apparent difference in glucose removal following exogenous insulin administration (Figure 1F, inset). HFD-fed rats had significantly higher serum insulin and plasma triaglyceride concentrations, compared to SD, whilst cholesterol levels were similar between the groups (Figure 1G-I and Table 1).

**Pressure myography**

Blood vessel contraction and relaxation were assessed in pressurized arteries using the open source pressure myograph system, VasoTracker[1]. Briefly, isolated mesenteric arteries were cleaned of adherent fat and mounted on resistance-matched glass pipettes in an arteriograph chamber. Arteries were pressurized to 70 mmHg (no flow) using a dual-reservoir pressure column system and left to equilibrate for at least 30 min in PSS (at 37ºC). Intraluminal flow (~200 µl min$^{-1}$) was established by a hydrostatic pressure gradient (10 cm $H_2O$) across the artery. Outer vessel diameter was continuously monitored with a CMOS camera and VasoTracker diameter tracking software. Phenylephrine (PE) was added to the physiological saline solution (PSS), which was continuously recirculated through the myograph chamber. The phenylephrine concentration was titrated (300 nM – 1 µM) to generate a 20% reduction in artery diameter in each experiment. Acetylcholine (ACh), $N^G$-nitro-L-arginine methyl ester (L-NAME), TRAM34, apamin, and 4-DAMP were each applied intraluminally. At the conclusion of each experiment, arteries were challenged with the NO donor, sodium nitroprusside (SNP, 10 µM). Contraction data are represented as the percent reduction from resting diameter. Relaxation data (from constricted diameter) are represented as the percent of maximal relaxation (constricted diameter to resting diameter).

**Preparation of arteries for endothelial cell imaging**

Arteries were cut open longitudinally, pinned *en face* (endothelial side up) to either: (1) the bottom of a Sylgard-coated flow chamber[2], for imaging with an upright microscope, or (2) on Sylgard blocks, for imaging on an inverted microscope [3]. The endothelium was then preferentially loaded with the $Ca^{2+}$ indicator, Cal-520/AM (5 µM; in DMSO with 0.02% Pluronic F-127) at 37°C for 30 minutes. In some experiments, inositol trisphosphate ($IP_3$) receptors ($IP_3Rs$) were directly activated by photolysing a caged form of $IP_3$. In these experiments, a membrane-permeant, photolabile version of the inositide (caged $IP_3$; caged IP3,4,5-dimethoxy-2-nitrobenzyl; 5 µM) with Pluronic F-127 (0.02%) was included in the $Ca^{2+}$-indicator solution[4, 5].

**Vascular reactivity and endothelial $Ca^{2+}$ levels in *en face* arteries**

Vascular reactivity and basal endothelial $Ca^{2+}$ levels were recorded by imaging Cal-520/AM fluorescence, excited at 488 nM using an LED illumination system (PE300Ultra, CoolLED, Andover, UK), on an upright fluorescence microscope (FN-1; Nikon, Tokyo, Japan). To maximize the field-of-view (~0.8 $\mu m^2$), this microscope was equipped with a 16X objective lens (0.8 NA; Nikon, Tokyo, Japan) and large-format (1024 x 1024 13 µm pixels) back-illuminated electron-multiplying charge-coupled device (EMCCD) camera (iXon 888; Andor, Belfast, UK). Illumination and image acquisition were controlled by uManager software[6]. Only arteries that, when cut open, fitted within the field-of-view (~250 µm diameter) were used. Arteries were perfused with PSS (1.5 ml $min^{-1}$) using a peristaltic pump. Contraction and dilation responses were assessed before and after incubation with L-NAME or BAPTA/AM (0.02% Pluronic F-127 for 30 minutes at 37°C). L-NAME remained present in the perfusion solution after incubation, whilst BAPTA/AM was washed out of the bath chamber at the end of the incubation.

Vascular reactivity was monitored using edge-detection algorithms[1, 2, 7]. PE-induced contraction was expressed as the percent reduction from resting diameter. ACh-evoked dilation (from constricted diameter) was expressed as a percentage of maximal relaxation (constricted diameter to resting diameter). Tissue movement resulting from contraction and dilation precluded an assessment of time-dependent changes in endothelial $Ca^{2+}$ levels. Instead, we measured endothelial $Ca^{2+}$ levels (expressed as arbitrary fluorescence units, A.U.) under resting conditions (in the absence of any pharmacological agent), or fractional changes in endothelial $Ca^{2+}$ levels preceding any tissue movement.

**High-resolution single-photon calcium imaging**

Endothelial cell activity was recorded by imaging Cal-520/AM fluorescence changes under either an inverted (high magnification/high-resolution; TE2000U; Nikon, Tokyo, Japan) or upright (low-magnification/high resolution; FN-1; Nikon, Tokyo, Japan) fluorescence microscope (TE2000U or FN-1; Nikon, Tokyo, Japan). The upright microscope system was the same as described above. The inverted microscope was equipped with a 100X objective lens (1.4 NA; Nikon, Tokyo, Japan), and the same LED illumination system and EMCCD as described above. Images were acquired at either 10 Hz (agonist-evoked activity) or 20 Hz (spontaneous activity).

**Analysis of stimulus-evoked calcium activity**

To assess the concentration-dependence of evoked $Ca^{2+}$ activity, we activated endothelial cells with increasing concentrations of ACh whilst recording $Ca^{2+}$ activity. The same field of endothelial cells, from the same artery, was visualized for each concentration, and there was a 10-minute wash/rest period between each stimulation. To identify the signaling pathways involved, responses to ACh (100 nM) were measured before and after treatment with various pharmacological agents (30-minute incubation, concentrations described in the text/figure legends) in the same preparation. In other experiments, $Ca^{2+}$ responses were evoked by photolysis of caged $IP_3$ using a xenon flashlamp (Rapp Optoelektronic, Hamburg, Germany). Light from the flashlamp, in the absence of caged $IP_3$, evoked no detectable $Ca^{2+}$ change in the endothelial cells.

Single-cell $Ca^{2+}$ activity was assessed as previously described[2, 8]. In brief, we first grouped imaging datasets by experiment and defined cellular regions of interest (ROIs) semi-automatically for each imaging session. We then defined ROIs for every cell within one of the datasets from a single experiment and projected these ROIs across all other datasets of the corresponding imaging session. Cells that were not within the field-of-view of each dataset in a group were excluded from analysis. We extracted $Ca^{2+}$ signals for each cell by averaging the fluorescence intensity across all pixels in each ROI. Traces were smoothed using a Savitzsly-Golay (21 point, 3$^{rd}$-order) filter, expressed as fractional changes in fluorescence ($F/F_0$) from baseline ($F_0$), and the discrete derivative was then calculated ($d(F/F_0)/dt$). The baseline was automatically determined by averaging the fluorescence intensity of the 100-frame portion of each trace that exhibited the least noise[2, 7].

For each endothelial cell, we automatically determined $Ca^{2+}$ activity using a peak-detection algorithm based on the discrete derivative. A $Ca^{2+}$ event was defined as an increase in the discrete derivative rising at least 10 standard deviations of baseline noise. Activity levels were quantified using the frequency of $Ca^{2+}$ spikes, the amplitude of $Ca^{2+}$ spikes ($\Delta F/F_0$; extracted from the corresponding $F/F_0$ trace), or steady-state (average) $Ca^{2+}$ levels ($\Delta F/F_0$) in each cell. Steady-state $Ca^{2+}$ levels were calculated by averaging $Ca^{2+}$ signals across a 60-second period immediately following the onset of spiking $Ca^{2+}$ activity. Where a cell exhibited no spiking activity, fluorescence intensity was averaged starting at the mean onset of activity in all other cells. Concentration-response data were normalized to the $Ca^{2+}$ rise induced by ionomycin (1 μM), applied at the end of each experiment.

**Assessment of endothelial heterogeneity**

To assess endothelial cell heterogeneity we constructed graph theoretical representations of the endothelial network using the Python programming language. To do this, we first generated reliable endothelial cell outlines using the "dilate no merge" plugin in FIJI[9, 10] to automatically identify neighboring endothelial cells. Then, we constructed binary adjacency matrices (A) to describe endothelial cell connectivity. Each element in the matrix (Aij) was set equal to a value of one if cells i and j were direct neighbors or zero otherwise. We disallowed self-connectivity by setting diagonal elements (Aii) to zero. These structural networks were represented as graphs using the NetworkX library[11] to reveal adjacency.

To assess cell sensitivity, the endothelium was stimulated with a maximal concentration of ACh and the cells that were unambiguously sensitive to ACh identified. A cell was considered ACh-sensitive if it responded before all other cells to which it was directly connected. To assess endothelial cell clustering, we identified the first 10% of cells to respond to a given stimulus. These fast responding population of cells are likely to be directly activated by the agonist rather than indirectly activated by signals arising in neighboring cells.

**Assessment of endothelial network communication**

To assess the extent of endothelial communication, we implemented a Python routine to calculate pairwise cross-correlation between neighboring endothelial cells[10, 12, 13]. To do this, discrete derivative $Ca^{2+}$ traces were each converted to single bit traces, assigning all time-points in the differentiated signals above threshold (10-fold standard deviation of baseline noise) to 1 and all other points to zero. Since cross-correlation coefficients between any cell pair could vary depending on the length or the exact portion of recording, we used a sliding window correlation analysis[14]. In particular, we calculated the mean cross-correlation coefficient between all pairs of neighboring cells in a 30-second interval beginning at the onset of stimulated $Ca^{2+}$ activity. We then shifted this interval throughout the time series with a step of 15 seconds, and for each step we again calculated the mean cross-correlation coefficient. For each neighboring cell pair, we used the grand mean of these cross correlation coefficients as a measure of coupling between cells. To determine if observed cross-correlation coefficients exceeded those expected by random chance, we used a permutation analysis. For each cell pair, the time course of one cell was shifted a random amount (holding individual cell activity levels constant) and coupling was again calculated for all cell pairs using these surrogate datasets. We repeated this step 1000 times for each dataset, creating a distribution of similarity indices expected at chance level for each cell pair.

**Assessment of coupling between spontaneous calcium events and myoendothelial projections**

To assess coupling between endothelial $Ca^{2+}$ events and smooth muscle cells (via myoendothelial projections, MEPs), spontaneous endothelial cell $Ca^{2+}$ activity and images of the underlying internal elastic lamina at the same site were recorded. Sites of initiation of spontaneous $Ca^{2+}$ events were identified automatically[4] and the distance from each site to the nearest hole in the internal elastic lamina (IEL) measured. To determine whether $Ca^{2+}$ event initiation sites preferentially occurred at MEPs, we used a permutation method to randomize the position of initiation sites. First, we generated surrogate datasets by randomly repositioning each initiation site to a random pixel location within the field-of-view, leaving the location of IEL unchanged. We then measured the distance between each repositioned initiation site and the closest IEL hole. For each dataset, we repeated this process 1000 times, creating a distribution of initiation-site to nearest IEL hole distance. To compare coupling of $Ca^{2+}$-events and MEP across experimental groups, we calculated the difference between the real $Ca^{2+}$-event-IEL hole separation and the corresponding separation in the randomized data, and expressed this difference as a fraction of the random separation. Resulting coupling coefficient values of one, indicates complete coupling ($Ca^{2+}$ events and IEL holes occur at precisely the same location), zero indicates random coupling ($Ca^{2+}$ events occur at random locations with respect to IEL holes), and a value less than zero, decoupling ($Ca^{2+}$ events do not occur at locations close to IEL holes).

### Statistics and data analysis

All image analysis was performed in FIJI[15] or using custom software written in Python 2.7. For food, water and energy measurements, intake per rat was calculated by dividing the total intake per cage by the number of animals in that cage and the reported n number is the number of cages. For all other experiments, the reported n represents the number of biological replicates (number of animals). In vascular reactivity experiments, a single artery was used for each biological replicate. In concentration-response $Ca^{2+}$ imaging studies, a single field of endothelial cells, from a single artery was used for each biological replicate. In experiments examining the spatial overlap of basal $Ca^{2+}$ events and IEL holes, results from three distinct fields of endothelial cells, from a single artery, were averaged for each biological replicate. Summary data are presented in text as mean ± standard error of the mean (SEM), and graphically as mean ± SEM (time-course data) or individual data points with the mean indicated. Paired data points in plots are indicated by connecting lines. Concentration-response data were computed according to a three-parameter dose–response model and compared using two-way ANOVA with Tukey's post-hoc test. All other data were analyzed using paired t tests, independent 2-sample t tests (with Welch's correction as appropriate), ordinary two-way ANOVA with multiple comparisons, or repeated measures (RM) two-way ANOVA with multiple comparisons as indicated in the respective figure or table legend. All statistical tests were two-sided. A p value of $< 0.05$ was considered statistically significant.

### Drugs and Solutions

L-NAME, TRAM-34, U73122 and U73343 were obtained from Tocris (St Louis, MO, USA). Cal-520/AM was from Abcam (Cambridge, MA, USA). Caged-$IP_3$ (caged-$IP_3$ 4,5–dimethoxy-2-nitrobenzyl) was obtained from Sichem (Germany). Pluronic F-127 was from Invitrogen (Carlsbad, CA, USA). All other drugs and chemicals were obtained from Sigma (St Louis, MO, USA). For the ITT, insulin was first prepared as a 25 U $ml^{-1}$ stock solution in 0.02 M HCl, and was then diluted to a 1 U $ml^{-1}$ working solution with distilled water. 2-aminoethoxydiphenyl borate (2-APB), 4-DAMP, BAPTA/AM, L-NAME, TRAM-34, U73122, U73343 were dissolved in DMSO and diluted to working concentration in PSS such that the total volume of DMSO was less than or equal to 0.1%. All other drugs were dissolved in water. The PSS consisted of (mM): 145 NaCl, 4.7 KCl, 2.0 MOPS, 1.2 $NaH_2PO_4$, 5.0 glucose, 0.02 EDTA, 1.17 $MgCl_2$, 2.0 $CaCl_2$, (adjusted to pH 7.4 with NaOH). In experiments using a $Ca^{2+}$-free PSS, $Ca^{2+}$ was substituted with $Mg^{2+}$ on an equimolar basis, and EGTA (1 mM) was included. Solutions were freshly prepared each day.

### Supplemental Figures and Figure Legends

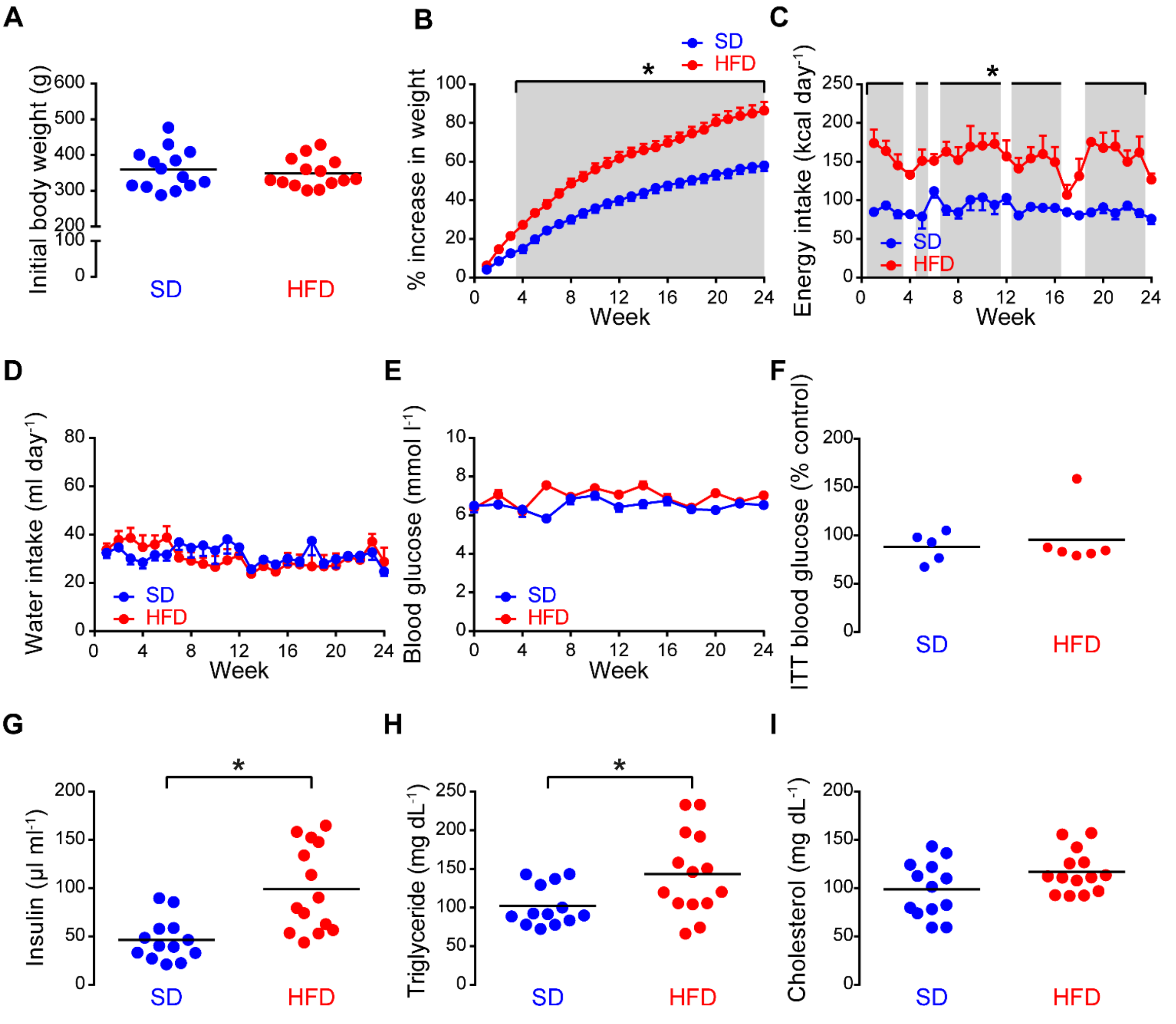

**Figure S1 – A high-fat diet induces prediabetic obesity in Wistar rats.** A) Initial body weight of rats fed the standard diet (SD) or the high-fat diet (HFD). Black line indicates the mean. B-F) Percentage weight gain (from initial body weight, B), mean energy intake (C), water intake (D), and blood glucose levels measured at rest (E) and 60 min after glucose loading (F, black line indicates the mean) for rats fed the standard (blue, n = 13) or high-fat (red, n = 14) diets. G-I) Serum insulin (G), triglyceride (H), and cholesterol in SD and HFD groups. Black line indicates the mean. * indicates $p < 0.05$ using repeated-measures two-way ANOVA with Sidak's multiple comparison test or unpaired t-test with Welch's correction, as appropriate. Data tabulated in Table S1.

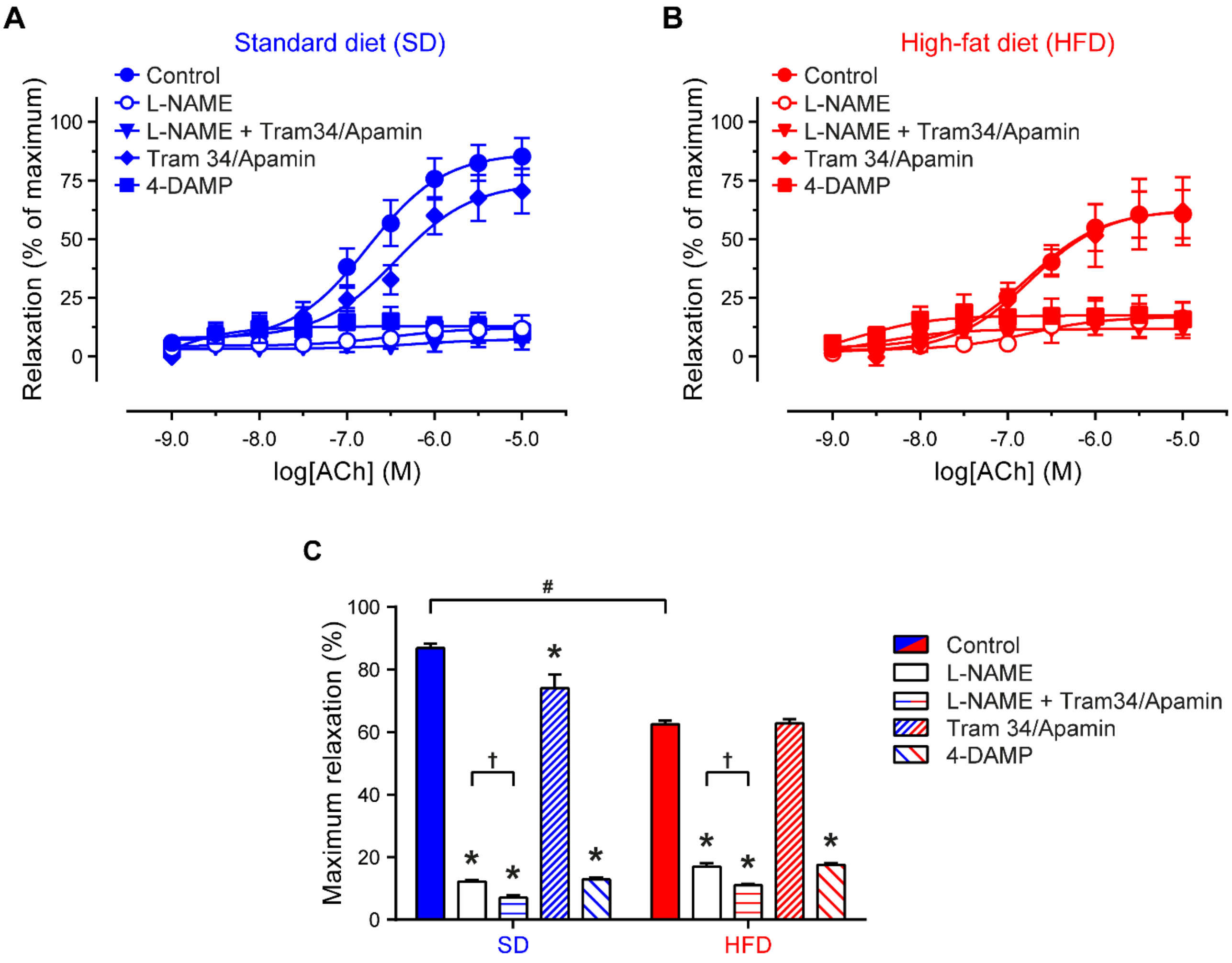

**Figure S2 – Effect of various inhibitors on ACh-induced vasodilation.** A-B) Summary of diameter data comparing the concentration-dependent responses of arteries, from rats fed either the standard diet (SD, panel A) or the high-fat diet (HFD, panel B), to intraluminal ACh. The concentration of ACh was increased cumulatively. All inhibitors were applied intraluminally at the following concentrations: L-NAME (100 µM), TRAM34 (1 µM), apamin (100 nM), 4-DAMP (1 µM). Curves are three-parameter sigmoid models. C) Summary of maximal relaxations to ACh (top of concentration-response curve). Data are given as mean ± SEM (n = 5 to 9). All data compared using two-way ANOVA with Tukey's post hoc test. *$p < 0.05$ versus corresponding control, # $p < 0.05$ vs SD control, † $p < 0.05$ versus corresponding L-NAME. Data corresponds to that shown in Figure 2 and tabulated in Table S2.

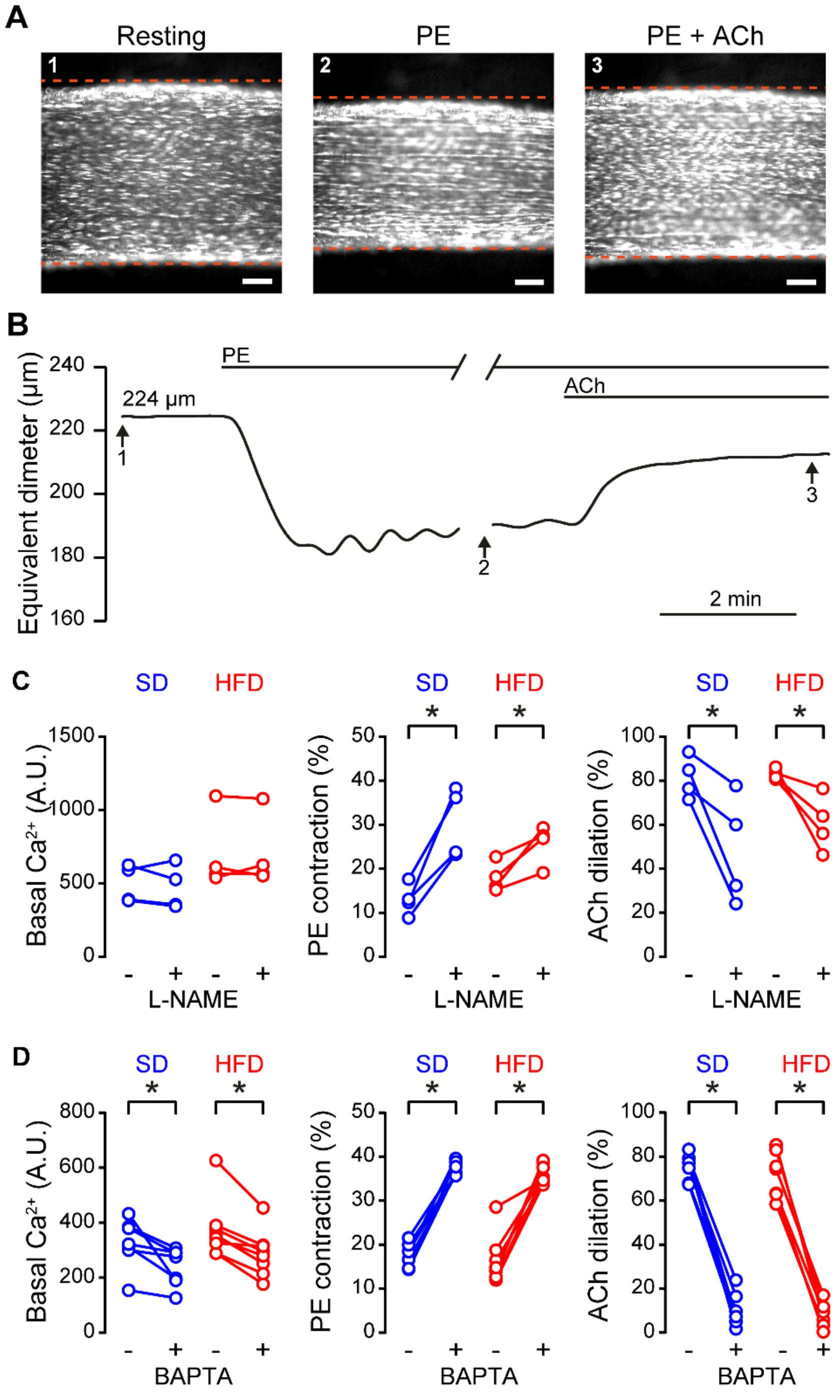

**Figure S3 –Endothelium-dependent vasodilation to ACh requires nitric oxide synthase and $Ca^{2+}$.** A-B) Fluorescence images showing an *en face* artery under resting conditions (1), after smooth muscle stimulation with PE (1 µM, 2), and then after stimulation of endothelial $Ca^{2+}$ using ACh (100 nM, 3). Vascular reactivity was assessed by tracking the edges of the artery. Scale bars = 100 µm. B) Equivalent diameter traces showing the full time-course of contraction and relaxation for the data shown in A. C) $Ca^{2+}$ levels (arbitrary fluorescence units, A.U.), PE-induced contraction, and ACh-induced dilation before and after nitric oxide synthase inhibition using L-NAME (100 µM, n = 4 for each group). D) $Ca^{2+}$ levels (arbitrary fluorescence units, A.U.), PE-induced contraction, and ACh-induced dilation before and after buffering endothelial $Ca^{2+}$ using BAPTA/AM (30 µM, n = 7 for SD, n = 8 for HFD). * $p < 0.05$, using paired t-test. Data tabulated in Tables S3-4.

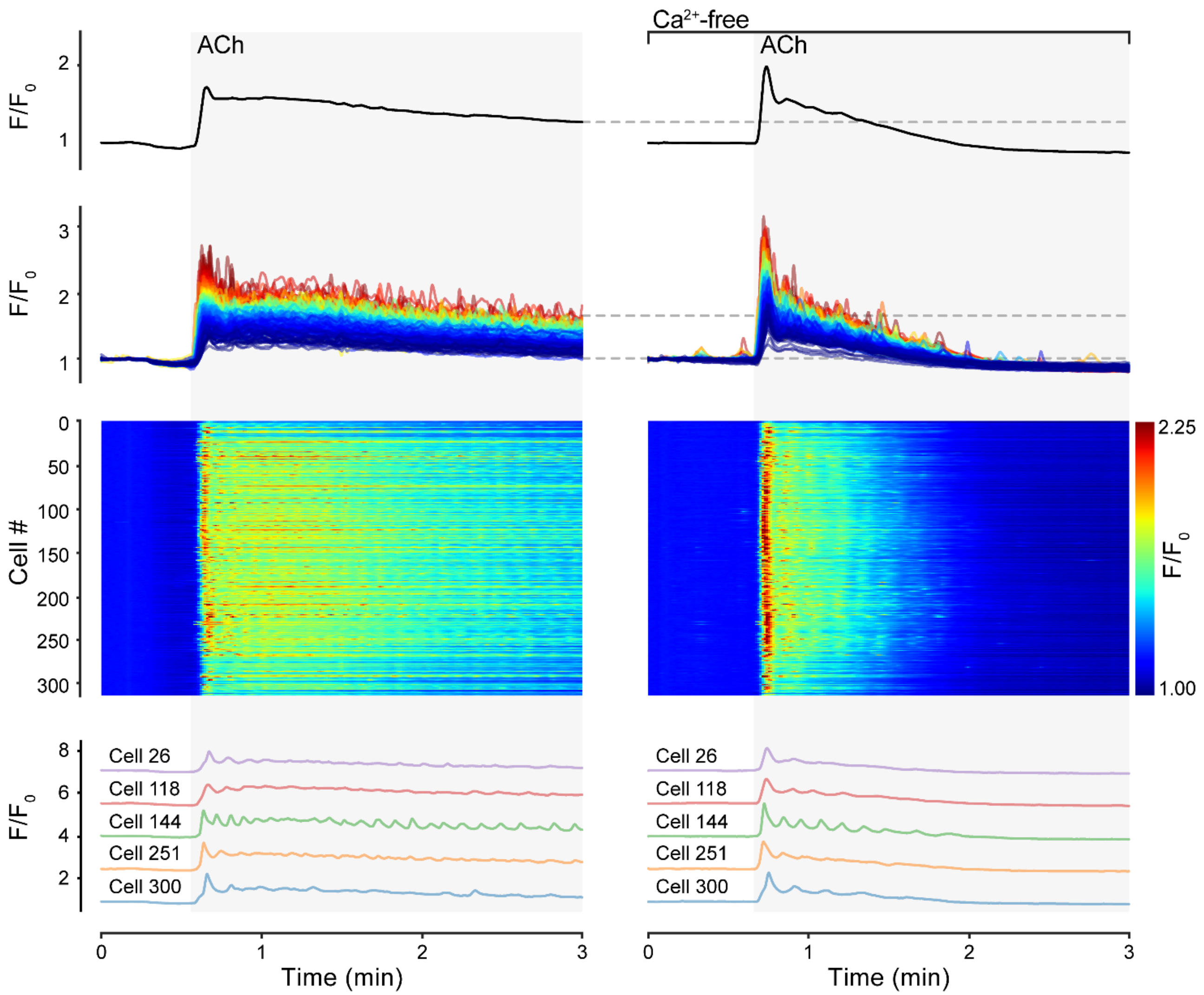

**Figure S4 – Acetylcholine induces $Ca^{2+}$ release in mesenteric endothelial cells.** The top row shows the average $Ca^{2+}$ signal from a field of ~300 mesenteric artery endothelial cells stimulated by ACh (100 nM) before (left) and after (right) the removal of external $Ca^{2+}$. $Ca^{2+}$-free PSS contained 1 mM EGTA. The second row shows $Ca^{2+}$ signals extracted from each cell in the field-of-view. $Ca^{2+}$ traces are color coded according to the amplitude of the response in the presence of external $Ca^{2+}$. The third row shows heatmaps indicating spiking $Ca^{2+}$ activity by a color scale. The bottom row shows example traces from five separate cells. Data corresponds to that shown in Figure 5 and tabulated in Table S7.

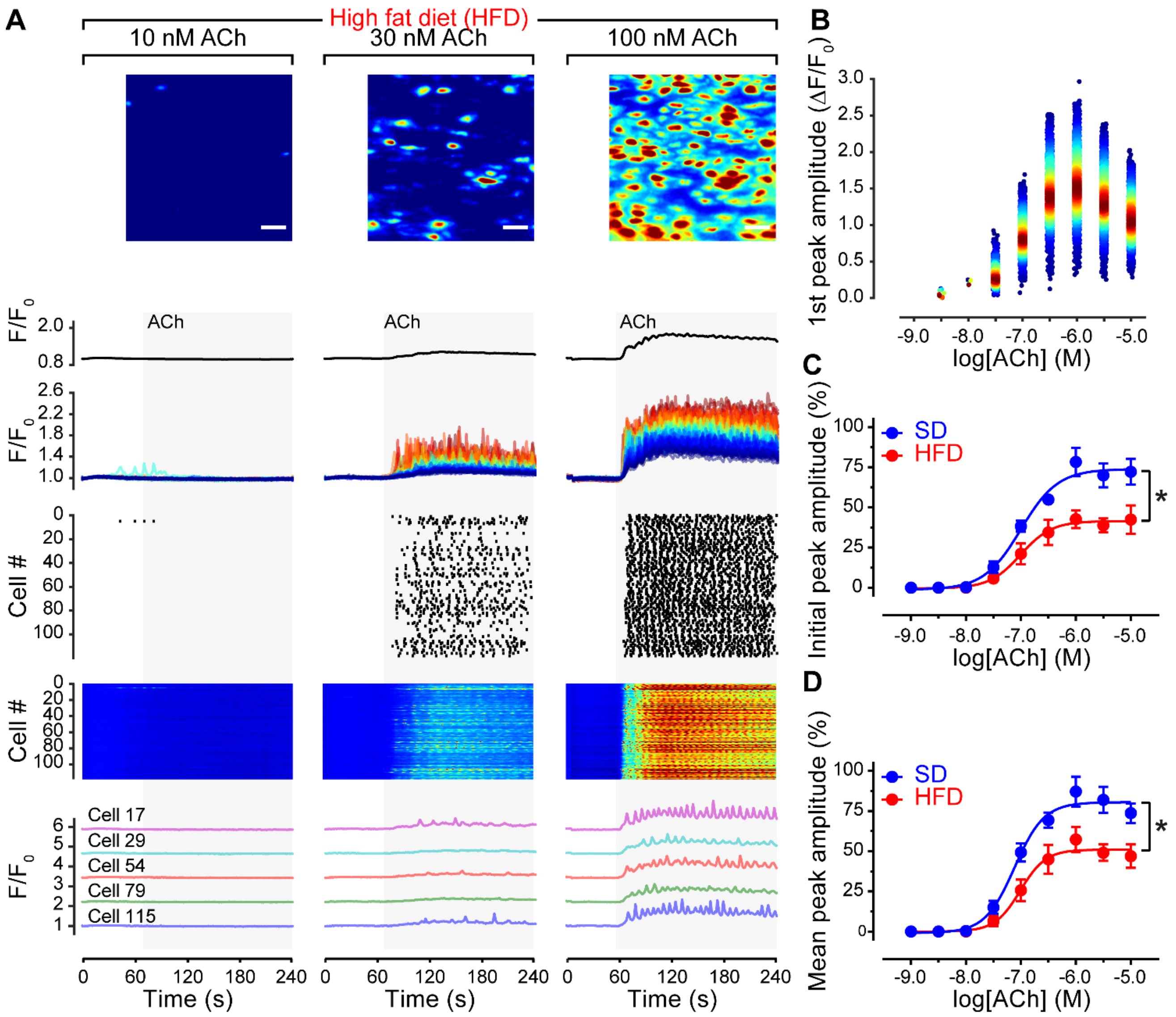

**Figure S5 – A high-fat diet impairs endothelial $Ca^{2+}$ signaling.** A) Concentration-dependence of ACh-evoked endothelial cell $Ca^{2+}$ activity. The top row displays pseudocolored $\Delta F/F_0$ maximum intensity projections of a single field of mesenteric endothelial cells (high-fat diet group, HFD) stimulated with 10 nM (left), 30 nM (middle) and 100 nM (right) ACh. Scale bars = 20 µm. The second row displays the average $Ca^{2+}$ signal across the field of view, whilst the third row displays $Ca^{2+}$ signals extracted from each cell in the field-of-view. $Ca^{2+}$ traces are color coded according to the amplitude of the response to 100 nM ACh. The fourth and fifth rows are rastergrams (fourth row) and heatmaps (fifth row), each indicating spiking $Ca^{2+}$ activity. The bottom row shows example traces from five separate cells. B) Concentration dependent activation of endothelial $Ca^{2+}$ signaling, as illustrated for each peak in $Ca^{2+}$ exhibited by each cell, for the full concentration-response experiment illustrated in panel A. C-D) Summary of $Ca^{2+}$ imaging data illustrating the concentration-dependence of the initial (C) and all (D) peaks in $Ca^{2+}$, evoked by ACh, in endothelium of rats fed the standard diet (SD, blue) and the high-fat diet (HFD, red). Data are mean ± SEM (n = 5 per group). * indicates significance ($p < 0.05$) by comparison of three-parameter fit parameters (top of sigmoid, log $EC_{50}$) using using two-way ANOVA with Sidak's multiple comparison test. Additional data tabulated in Table S5.

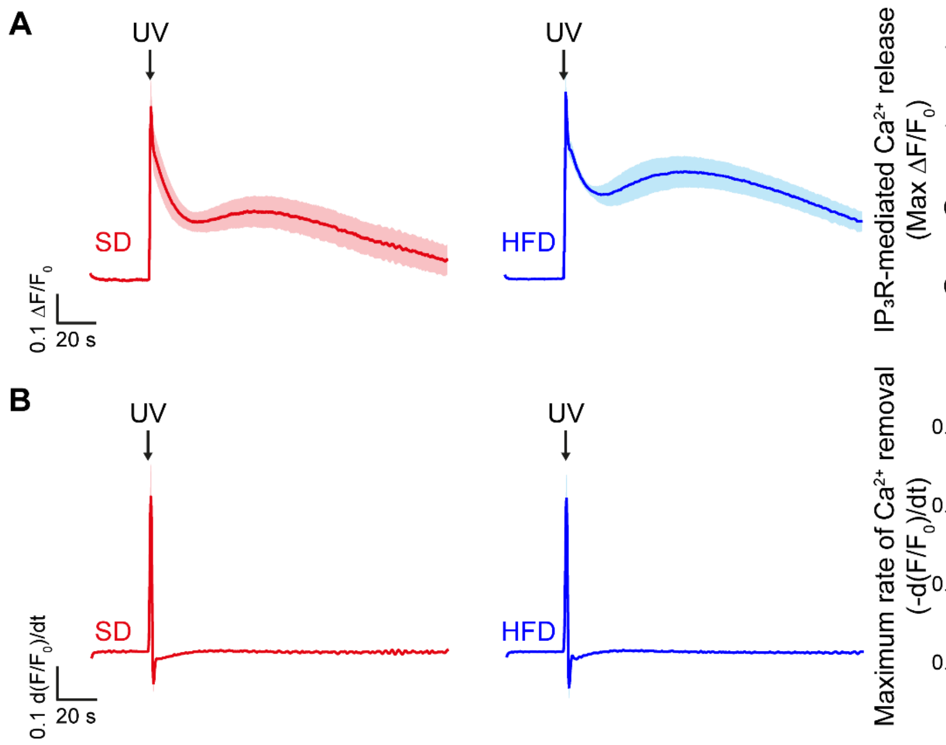

**Figure S6 – $IP_3$-mediated $Ca^{2+}$ release, and $Ca^{2+}$ removal is unaltered in obesity.** A) $Ca^{2+}$ responses to photolysis of caged $IP_3$ in mesenteric artery endothelial cells from rats fed a standard diet (SD, left) or a high-fat diet (HFD, middle). Each trace is the mean ± S.E.M of the global endothelial $Ca^{2+}$ response from 5 (SD) or 6 HFD) biological replicates. The right panel is summary data indicating the ability of $IP_3$ to evoke $Ca^{2+}$ release (max $\Delta F/F_0$ of response to photolysis of caged $IP_3$). B) Differentiated $Ca^{2+}$ responses to photolysis of caged $IP_3$ (original $F/F_0$ data shown in A), showing the similar rate of change of intracellular $Ca^{2+}$ concentration in the endothelium of the SD (left) and HFD (right) groups. The right panel is summary data indicating the maximum rate of $Ca^{2+}$ removal (minima of derivate $Ca^{2+}$ trace) in each group.

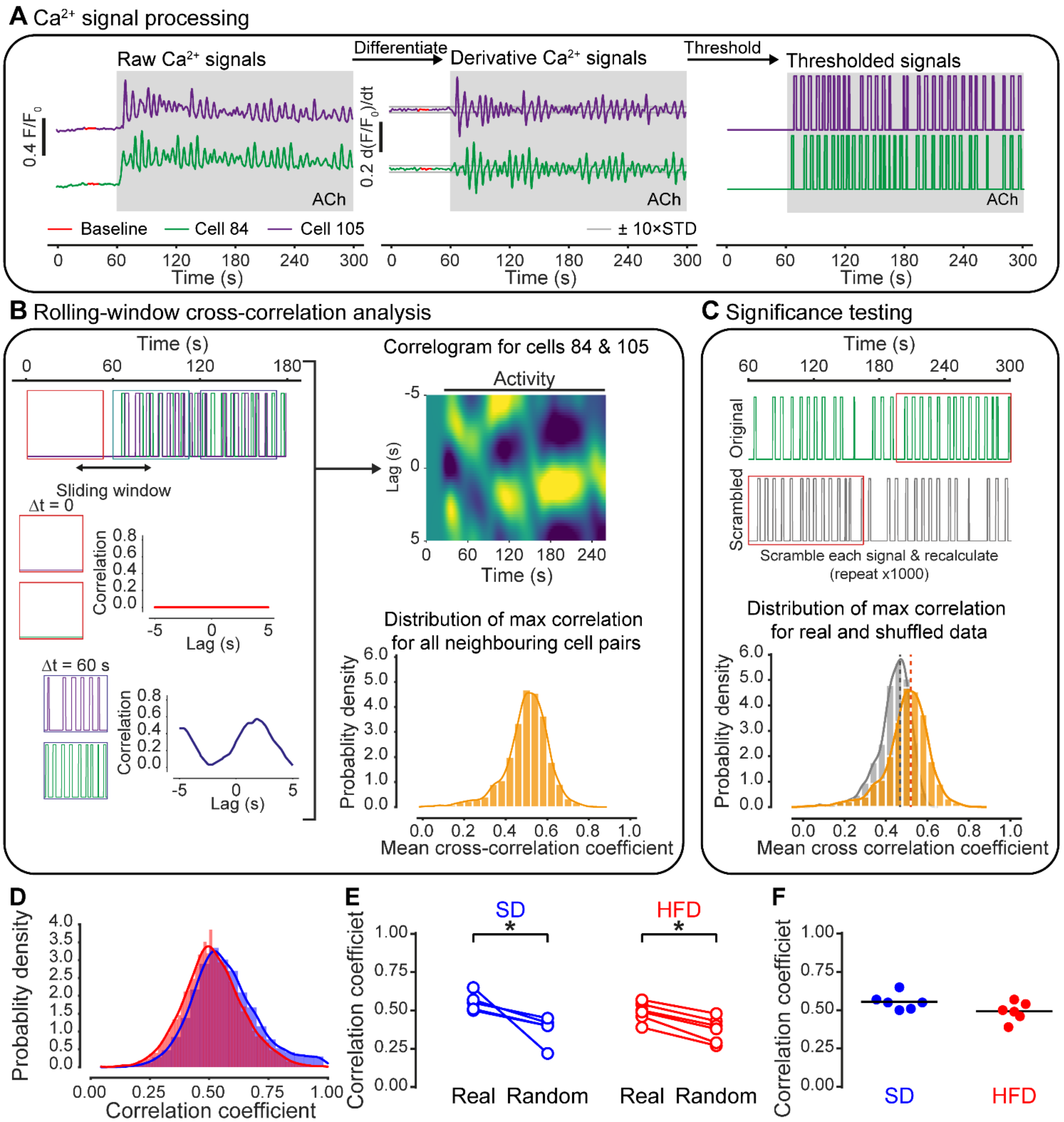

**Figure S7 –Functional connectivity is unaltered in obesity.** A-C) Methods for estimating functional connectivity from $Ca^{2+}$ imaging data. A) Processing of $Ca^{2+}$ signals for correlation analysis. False correlations between $Ca^{2+}$ signals due to underlying trends, were eliminated by taking the first derivative (middle). This process is carried out for every cell identified when $Ca^{2+}$ signals occurred. Artefacts due to amplitude fluctuations and sparse data (zero values) were eliminated by normalizing the signal magnitude between zero and one (thresholded; bottom) and Gaussian noise added to prevent correlations arising between signals with low activity. (**C**) Trial shuffling and scrambling signals were used to generate artificial randomized data for permutation testing. The thresholded data (left) was first deranged (randomly shuffled; middle) and further randomized by shuffling each signal (right) from a random time point. (**D**,**E**) Correlation as a function of distance between cells for original (D) and randomized (E) $Ca^{2+}$ signals from an experiment imaging ~150 cells. Repeating the randomized analysis (thousands of times) permits permutation testing. Comparing correlation values against the randomized distribution enabling significant correlations to be identified. (**F–I**) $Ca^{2+}$ image overlaid with: (F) functional network map showing all possible connections between cells; (G) structural network map showing all physical connections (e.g., gap junctions); (H) refined functional network map with significant correlations obtained by permutation testing and refined using the structural network; (**H**) colour-coded cell ensembles identified from $Ca^{2+}$ imaging data. * indicates $p < 0.05$ using paired (E) or unpaired (F) t-test.

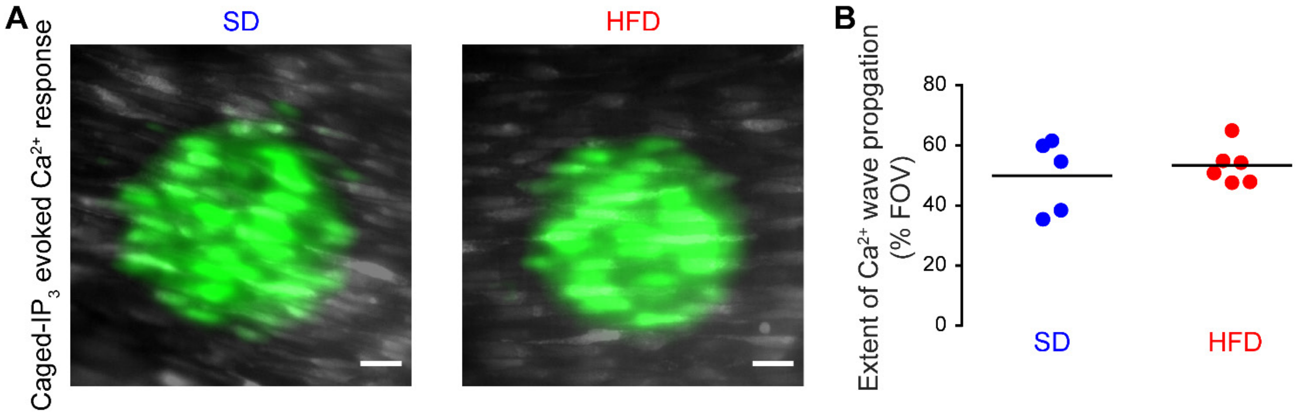

**Figure S8 – $Ca^{2+}$ wave propagation is unaltered in obesity.** A) Composite $Ca^{2+}$ images illustrating $Ca^{2+}$ activity in response to local photolysis of caged $IP_3$ in the endothelium of rats fed a standard diet (SD, left) or a high-fat diet (HFD, right). Scale bars = 20 µm B). Summary data illustrating the extent of $Ca^{2+}$-wave propagation. Statistical comparisons ($p < 0.05$ considered significant) made using unpaired t-tests with Welch's correction.

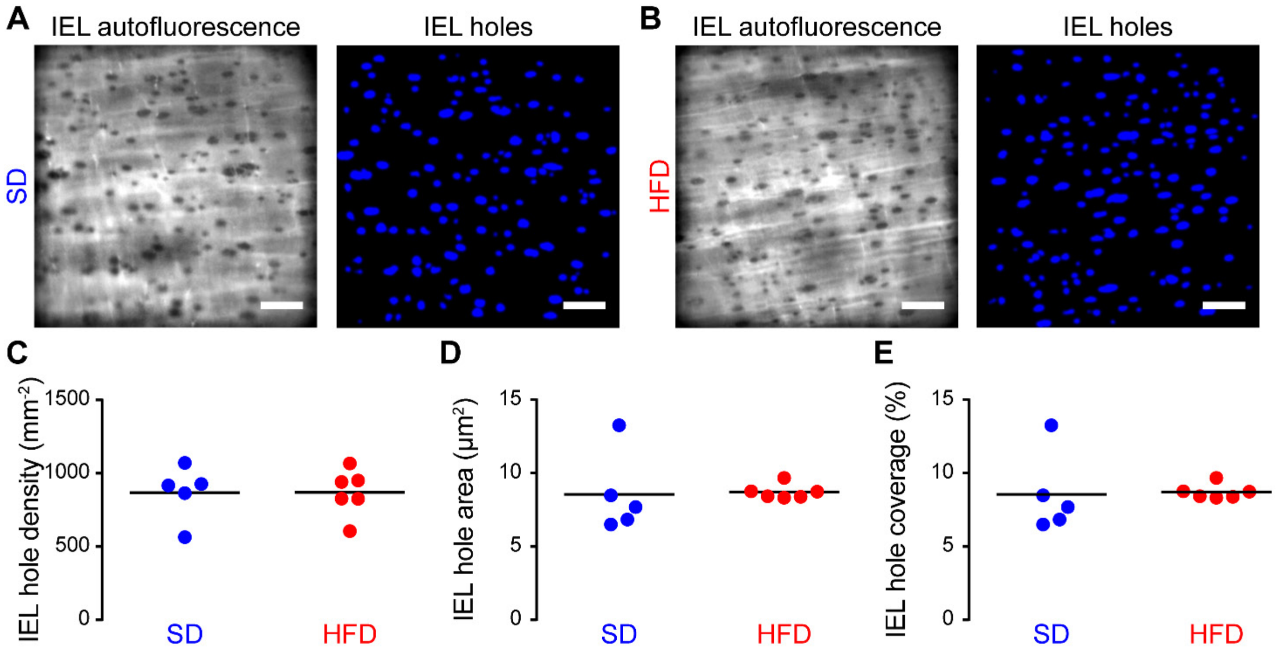

**Figure S9 – IEL structure is unaltered in obesity.** A-B) Representative images of IEL holes in mesenteric arteries from rats fed either the standard diet (SD, A) or the high-fat diet (HFD, B) diet. Left panels show raw elastin autofluorescence images. These images were processed and inverted to highlight IEL holes (right panels). Scale bars = 20 µm. C-E) Summary data (black line shows mean) showing the effect of diet IEL hole density (C), IEL hole area (D), and the area of IEL occupied by holes (E). Statistical comparisons ($p < 0.05$ considered significant) made using unpaired t-tests with Welch's correction. Data tabulated in Table S6.

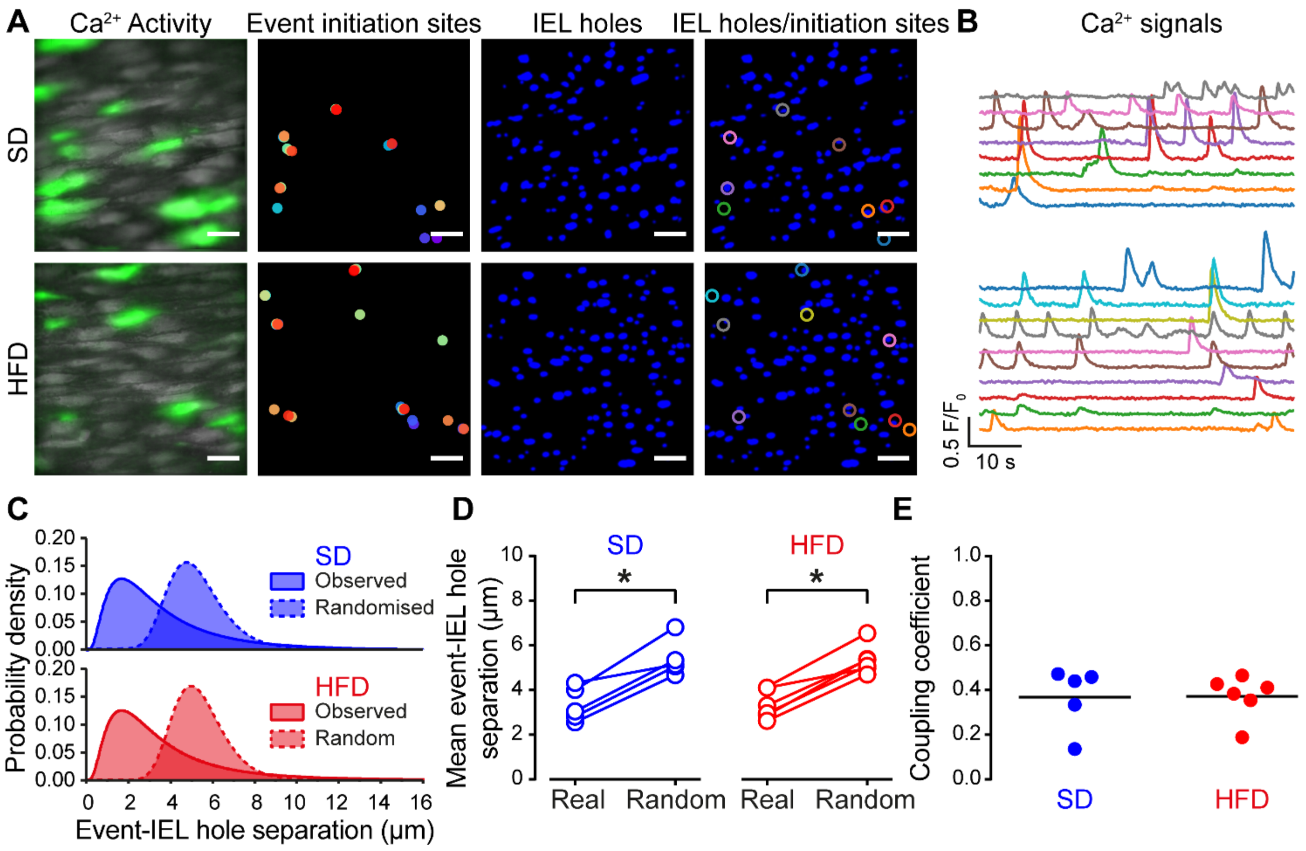

**Figure S10 – Myoendothelial coupling is unaltered in obesity.** A-B) Representative images of basal (unstimulated) endothelial $Ca^{2+}$ activity and the holes in the underlying IEL (IEL holes) in small mesenteric arteries from rats fed either the standard diet (SD) or the high-fat diet (HFD) diet. The first column shows averaged fluorescence intensity (grey) with $Ca^{2+}$ activity overlaid (green, 60 s activity). The second column shows the sites where basal $Ca^{2+}$ events initiated. The third column shows holes in the underlying IEL. The final column shows ROIs corresponding to $Ca^{2+}$ event initiation sites overlaid on the IEL hole image. Scale bars = 20 µm. B) Color-coded $Ca^{2+}$ traces extracted from the initiation sites indicated in A. C) Distributions (Gaussian-modified exponential) of the centroid-centroid distance between $Ca^{2+}$ event initiation sites and the nearest IEL hole for real (solid lines) and randomized (dashed lines) data. D) Paired plots showing the mean separation for real and random data. Each data point is the mean of at least three separate fields of endothelial cells from a single animal. E) Mean coupling coefficient of $Ca^{2+}$ events and IEL holes. The coupling coefficient was calculated for each individual dataset by expressing the difference between the real $Ca^{2+}$ event-IEL hole separation and the corresponding random separation, as a fraction of the random separation. Thus, a coupling coefficient of one or zero indicates complete coupling or no coupling, respectively. Black line indicates the mean. * indicates $p < 0.05$ using paired t-test. Data tabulated in Table S6.

### Supplemental Tables and supporting information

#### Table S1. General animal data

| | SD | HFD | Significance |
|---|---|---|---|
| Starting body weight, g | 358 ± 16 | 349 ± 11 | |
| Body weight at 24 weeks, g | 565 ± 26 | 654 ± 33 | |
| Increase in body weight, % | 58 ± 3 | 86 ± 4 | * |
| Food intake, g day$^{-1}$ | 28 ± 3 | 35 ± 3 | |
| Energy intake, kcal day$^{-1}$ | 89 ± 5 | 154 ± 11 | * |
| Water intake, ml day$^{-1}$ | 32 ± 2 | 31 ± 3 | |
| Blood glucose, mmol l$^{-1}$ | 6.5 ± 0.1 | 6.9 ± 0.1 | |
| Serum insulin, µl ml$^{-1}$ | 46 ± 6 | 99 ± 12 | * |
| Serum cholesterol, µg µl$^{-1}$ | 99 ± 8 | 117 ± 6 | |
| Serum triglyceride, nmol l$^{-1}$ | 102 ± 7 | 143 ± 14 | * |

Data are mean ± SEM. * $p < 0.05$, standard diet (SD) versus high-fat diet (HFD) using unpaired t-test with Welch's correction.

#### Table S2. Properties of mesenteric arteries from rats fed a standard diet (SD) or a high-fat diet (HFD).

| | SD | HFD |
|---|---|---|
| Outer diameter (unpressurised) | 266 ± 6 (8) | 285 ± 11 (8) |
| Outer diameter (70 mmHg) | 399 ± 10 (8) | 427 ± 17 (8) |
| Maximum relaxation to ACh | | |
| Control | 86.8 ± 1.2 (7) | 62.5 ± 1.1 (9) # |
| L-NAME (100 µM) | 12.2 ± 0.6 (7) * | 17.0 ± 1.2 (7) * |
| L-NAME + TRAM34 (1 µM) / Apamin (100 nM) | 7.4 ± 0.8 (5) * † | 11.7 ± 0.4 (7) * † |
| TRAM34 (1 µM) / Apamin (100 nM) | 74.0 ± 4.5 (6) * | 62.8 ± 1.3 (6) |
| 4-DAMP | 12.9 ± 0.7 (6) * | 17.5 ± 0.6 (6) * |
| Maximum relaxation to SNP (10 µM, % maximum) | 92.9 ± 2.9 (8) | 90.2 ± 4.0 (8) |

Data are mean ± SEM. Numbers in parenthesis indicate n-values (number of biological replicates). SD, standard diet HFD, high fat diet. *, $p < 0.05$ versus within group control (no treatment). †, $p < 0.05$ within group response in the presence of L-NAME; #, $p < 0.05$ versus SD control. Statistical comparisons made using two-way ANOVA with Tukey's multiple comparisons test.

**Table S3. Effect of L-NAME on basal endothelial $Ca^{2+}$ levels and vascular reactivity measured in the same *en face* arteries.**

| Group | | Control | L-NAME (100 µM) | Significance |
|---|---|---|---|---|
| SD (n = 4) | Basal $Ca^{2+}$ levels (A.U.) | 498 ± 64 | 472 ± 75 | |
| | PE Contraction | 13 ± 2 | 30 ± 4 | * |
| | ACh Relaxation | 81 ± 5 | 49 ± 12 | * |
| HFD (n = 4) | Basal $Ca^{2+}$ levels (A.U.) | 703 ± 132 | 706 ± 125 | |
| | PE Contraction | 18 ± 2 | 26 ± 2 | * |
| | ACh Relaxation | 84 ± 2 | 60 ± 6 | * |

Data are mean ± SEM, n-values are number of biological replicates. * $p < 0.05$, versus within group control using paired t test.

**Table S4. Effect of BAPTA on basal endothelial $Ca^{2+}$ levels and vascular reactivity measured in the same *en face* arteries.**

| Group | | Control | BAPTA (30 µM) | Significance |
|---|---|---|---|---|
| SD (n = 7) | Basal $Ca^{2+}$ levels (A.U.) | 327 ± 34 | 240 ± 26 | * |
| | PE Contraction | 18 ± 1 | 37 ± 1 | * |
| | ACh Relaxation | 76 ± 2 | 9 ± 3 | * |
| HFD (n = 7) | Basal $Ca^{2+}$ levels (A.U.) | 375 ± 38 | 288 ± 29 | * |
| | PE Contraction | 16 ± 2 | 36 ± 1 | * |
| | ACh Relaxation | 73 ± 4 | 4 ± 3 | * |

Data are mean ± SEM, n-values are number of biological replicates. * $p < 0.05$, versus within group control using paired t test.

**Table S5. Effect of pharmacological intervention on ACh-evoked endothelial $Ca^{2+}$ signaling metrics in rats fed a standard diet (SD).**

| Treatment | Cells responding (%) | $Ca^{2+}$ signaling parameter<br>Initial response (Peak $F/F_0$) | Steady-state response (Mean $F/F_0$) |
|---|---|---|---|
| 4-DAMP (n = 6) | 6 ± 1 * | 2 ± 1 * | 7 ± 2 * |
| $Ca^{2+}$-free (n = 6) | 98 ± 2 | 132 ± 14 | 66 ± 7 |
| U73343 (n = 7) | 73 ± 11 | 62 ± 15 * | 73 ± 12 |
| U73312 (n = 7) | 30 ± 12 * # | 19 ± 8 * # | 45 ± 11 * # |
| 2-APB (n = 5) | 14 ± 10 * | 14 ± 13 * | 53 ± 18 * |
| Caffeine (n = 5) | 41 ± 12 * | 15 ± 6 * | 22 ± 3 * |
| RuR (n = 5) | 108 ± 2 | 136 ± 19 | 152 ± 15 * |

Data are mean ± SEM, expressed as a percentage of the control response obtained prior to the pharmacological intervention; n-values are number of biological replicates. $Ca^{2+}$-free PSS contained 1 mM EGTA. Concentrations of pharmacological inhibitors were: 4-DAMP (1 μM); U73343 (2 μM); U73122 (2 μM), 2-APB (100 μM); caffeine (10 mM); RuR (5 μM). *, $p < 0.05$ versus within group control. #, $p < 0.05$ versus versus within-group U73343. Statistical comparisons made on raw data using repeated-measures two-way ANOVA with Tukey's or Dunnett's multiple comparison test, as appropriate.

**Table S6. Effect of pharmacological intervention on ACh-evoked endothelial $Ca^{2+}$ signaling metrics in rats fed a high-fat diet (HFD).**

| Treatment | Cells responding (%) | $Ca^{2+}$ signaling parameter<br>Initial response (Peak $F/F_0$) | Steady-state response (Mean $F/F_0$) |
|---|---|---|---|
| 4-DAMP (n = 7) | 9 ± 4 * | 2 ± 1 * | 25 ± 8 * |
| $Ca^{2+}$-free (n =7) | 100 ± 1 | 115 ± 9 | 80 ± 10 |
| U73343 (n = 7) | 92 ± 3 | 71 ± 6 * | 80 ± 9 |
| U73312 (n = 7) | 51 ± 12 * # | 29 ± 9 * # | 42 ± 8 * # |
| 2-APB (n = 5) | 27 ± 15 * | 4 ± 2 * | 36 ± 8 * |
| Caffeine (n = 7) | 39 ± 10 * | 2 ± 1 * | 30 ± 7 * |
| RuR (n = 5) | 117 ± 15 | 110 ± 17 | 108 ± 6 |

Data are mean ± SEM, expressed as a percentage of the control response obtained prior to the pharmacological intervention; n-values are number of biological replicates. $Ca^{2+}$-free PSS contained 1 mM EGTA. Concentrations of pharmacological inhibitors were: 4-DAMP (1 μM); U73343 (2 μM); U73122 (2 μM), 2-APB (100 μM); caffeine (10 mM); RuR (5 μM). *, $p < 0.05$ versus within group control control. #, $p < 0.05$ versus within-group U73343. Statistical comparisons made on raw data using repeated-measures two-way ANOVA with Tukey's or Dunnett's multiple comparison test, as appropriate.

**Table S7. Concentration-response curve parameters of ACh-evoked endothelial $Ca^{2+}$ signaling in mesenteric arteries from rats fed a standard diet (SD) or a high-fat diet (HFD).**

| $Ca^{2+}$ signaling metric | Curve parameter | SD (n = 5) | HFD (n = 6) | Significance |
|---|---|---|---|---|
| % cells responding | Top (%) | 100.0 ± 0.1 | 99.1 ± 1.5 | |
| | log $EC_{50}$ | -7.69 ± 0.01 | -7.47 ± 0.03 | |
| Oscillation frequency | Top (peaks $min^{-1}$) | 7.7 ± 0.2 | 6.9 ± 0.3 | |
| | log $EC_{50}$ | -7.531 ± 0.03 | -7.32 ± 0.07 | |
| Average $Ca^{2+}$ response | Top (% maximal) | 73.7 ± 3.6 | 46.1 ± 2.0 | * |
| | log $EC_{50}$ | -7.041 ± 0.09 | -7.0 ± 0.07 | |
| Amplitude of first $Ca^{2+}$ peak | Top (% maximal) | 76.7 ± 3.4 | 43.8 ± 1.9 | * |
| | log $EC_{50}$ | -6.94 ± 0.09 | -6.94 ± 0.09 | |
| Amplitude of all $Ca^{2+}$ peaks | Top (% maximal) | 84.7 ± 4.5 | 53.9 ± 3.6 | * |
| | log $EC_{50}$ | -7.05 ± 0.12 | -6.97 ± 0.14 | |

Data are mean ± SEM, n-values are number of biological replicates. Data modelled using three-parameter concentration response curve. * $p < 0.05$, standard diet (SD) versus high-fat diet (HFD) using two-way ANOVA with Sidak's multiple comparison test.

**Table S8. IEL structure and myoendothelial coupling in rats fed a standard or high-fat diet.**

| $Ca^{2+}$ signaling metric | | SD (n = 5) | HFD (n = 6) | Significance |
|---|---|---|---|---|
| IEL hole size ($\mu m^2$) | | 8.6 ± 1.2 | 8.7 ± 0.2 | |
| IEL hole density ($mm^{-2}$) | | 869 ± 83 | 870 ± 65 | |
| IEL hole coverage (% of IEL area) | | 6.8 ± 0.2 | 7.5 ± 0.5 | |
| Initiation site – IEL hole separation | Observed | 3.3 ± 0.3 | 3.3 ± 0.3 | * |
| | Random | 5.4 ± 0.4[#] | 5.3 ± 0.3[#] | |
| Coupling coefficient | | 0.37 ± 0.03 | 0.37 ± 0.04 | * |

Data are mean ± SEM, n-values are the number of biological replicates. # $p < 0.05$ versus within group control using paired. All other data analyzed using t-test with Welch's correction (* $p < 0.05$).
